# Macrophage-derived extracellular vesicles promote T cell–dependent inflammatory pain resolution

**DOI:** 10.64898/2026.03.03.706465

**Authors:** Jason R. Wickman, Richa Pande, Jason DaCunza, Deepa Reddy, Yuzhen Tian, Ezgi Ecem Kasimoglu, Roshell Muir, Elias K. Haddad, Seena K. Ajit

**Affiliations:** Department of Pharmacology & Physiology, Drexel University College of Medicine, Philadelphia, Pennsylvania 19102, USA; Microbiology and Immunology Graduate Program, Drexel University College of Medicine, Philadelphia, Pennsylvania 19102, USA; Division of Infectious Diseases and HIV Medicine, Department of Medicine, Drexel University College of Medicine, Philadelphia, Pennsylvania 19102, USA; Department of Microbiology and Immunology, Wake Forest University School of Medicine, Winston-Salem, NC 27101, USA

## Abstract

Inflammatory pain resolution is increasingly recognized as an active, immune-regulated process, yet the adaptive immune mechanisms that govern this process remain poorly defined. We previously demonstrated that intrathecal administration of macrophage-derived small extracellular vesicles (sEVs) from unstimulated (sEV) or LPS-stimulated (sEV^+^) RAW 264.7 cells accelerate resolution of complete Freund’s adjuvant (CFA)-induced inflammatory pain in male mice. However, the immunological mechanisms underlying this effect remain undefined. Given growing evidence that T cells regulate inflammatory pain resolution, we investigated whether macrophage-derived sEVs engage adaptive immune pathways to promote recovery. *In vitro*, both sEV and sEV^+^ enhanced T cell activation, with sEV^+^ exhibiting greater immunostimulatory capacity. Direct effects on T cells were modest; instead, sEV^+^ induced robust antigen-presenting cell (APC)-dependent T cell activation characterized by increased costimulatory molecule expression and enhanced Th1 polarization. Loss-of-function and rescue studies in *Rag2*^−/−^ mice demonstrate that T cells are required for late-phase sEV^+^-mediated inflammatory pain resolution. *In vivo*, sEV^+^ elicited immunostimulatory responses in intrathecal-draining cervical and CFA-draining sacral/internal iliac lymph nodes. Together, these findings identify adaptive immune engagement as a critical mediator of sEV^+^-induced pain resolution and position macrophage-derived sEVs as a cell-free immunotherapeutic modality that harnesses endogenous T cell–dependent mechanisms of active inflammatory pain resolution.

**Graphical Abstract:** 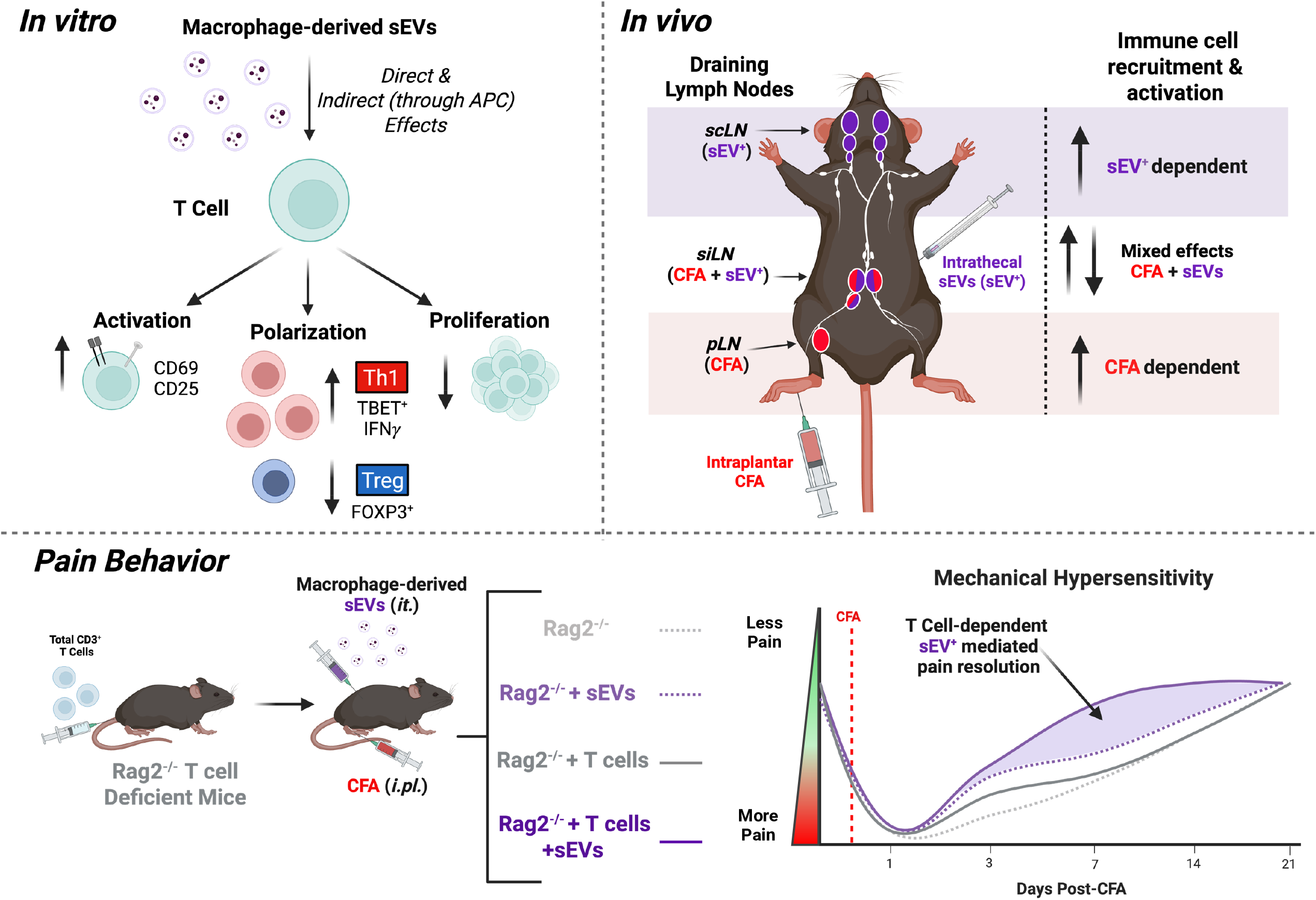

## Introduction

Small extracellular vesicles (sEVs) are lipid bilayer enclosed vesicles released naturally by cells with an average size range of 30 and 150 nm. sEVs contain proteins, lipids, and nucleic acids, and play an important role in neuroimmune crosstalk by facilitating transfer of complex cargo between cells[1, 2]. As sEV contents vary with the cell of origin and its functional state, the utility of sEVs as a potential therapeutic platform has garnered increasing interest across a wide range of diseases[3, 4].

Current therapeutic applications have strongly focused around sEVs from mesenchymal stem cells (MSCs) due to their low immunogenicity and regenerative properties[5]. However, in contrast, sEVs derived from immune cells have received interest due to their potential immunostimulatory functions, particularly in cancer therapies where boosting a directed immune response is desirable. sEVs derived from APCs such as dendritic cells and B cells have been shown to carry peptide-loaded MHC complexes (p-MHC) and co-stimulatory molecules such as CD80/CD86. These APC-derived sEVs can either directly present p-MHC complexes and costimulatory molecules to T cells, or indirectly where sEV p-MHC are processed by an APC or re-displayed on its surface[2, 6–8]. The immunostimulatory functions and therapeutic potential of macrophage-derived sEVs, particularly in their capacity to act as antigen-presenting cell–like mediators, remain less well characterized.

Our previous studies showed a single intrathecal injection of murine macrophage RAW 264.7 cell-derived sEVs promoted resolution of thermal hyperalgesia and mechanical allodynia[9], indicating the therapeutic potential of macrophage derived sEVs to promote pain resolution. Macrophages are an integral component of the innate immune system and depending on their activation, polarization, and local environmental cues they can accentuate damage or promote tissue repair[10, 11]. While neuronal mechanisms are critical for peripheral and central pain transmission recently contributions of aberrant neuroimmune signaling underlying the development of chronic pain are recognized[12–16].

Importantly, T cells have been shown to drive mechanisms that promote the resolution of inflammatory and neuropathic pain and are emerging as potential therapeutic target for chronic pain[17–19]. CD4^+^ T cells play an important role in endogenous analgesia mediated by enkephalins[20–22] and regulatory T cells (Treg) have been implicated in controlling pain hypersensitivity[23] and suppressing pro-inflammatory Th1 responses at the site of nerve injury[24]. Additionally, CD8^+^ T cells are important for the resolution of cisplatin-induced neuropathic pain and educating them prior to insult aided in pain resolution[25–27]. We had previously identified sEVs derived from LPS stimulated macrophages were enriched in RNA implicated in antigen processing, cross-presentation, and MHC-I presentation, suggesting the capacity to modulate T cell responses^29^. It is not known how intrathecal therapeutic administration of macrophage derived sEVs alters T cells *in vivo*.

We hypothesized that macrophage-derived sEVs activate T cells, and that T cell responses are required for late-phase sEV-mediated inflammatory pain resolution. We investigated whether the immunostimulatory effects of macrophage-derived sEVs on T cells *in vitro* are mediated through enhanced T cell activation, either directly, or indirectly via uptake by APCs, and examined whether sEV-mediated inflammatory pain resolution depends on T cell responses *in vivo*.

## Results

### Macrophage-derived (RAW 264.7) sEVs express myeloid and immune activation surface markers needed to directly activate T cells

sEVs were collected from the media of murine RAW 264.7 macrophage cells that were either untreated (sEV) or treated with 1 µg/mL LPS. Nanoparticle tracking analysis showed that sEVs had a particle size between 50-400 nm in size, with a mean size of 155 (sEV) and 173 (sEV^+^) nm, consistent with reported size ranges of sEVs **(Figure 1A)**. We have previously demonstrated the presence of sEV specific markers such as tetraspanin CD81 and ESCRT associated protein ALIX and absence of endoplasmic reticulum protein Calnexin in our EV preparations from RAW 264.7 cells[9]. We explored whether sEVs express immune surface markers associated with macrophage activation and antigen presentation, which would be necessary for sEVs to directly activate T cells. Surface staining of sEVs demonstrated the presence of tetraspanins known to be enriched in sEVs (CD9, CD63, CD81), myeloid markers (CD11b, CSF1R), and markers important for direct activation of T cells through antigen presentation and co-stimulation (CD40, MHCI, CD86) **(Figure 1B-C)**. We also investigated the presence of LPS as a contaminant in our sEV preparations. LPS was undetectable in sEV^+^ using a chromogenic endotoxin assay at a sensitivity of 0.01 EU/mL (data not shown).

**Figure 1.**
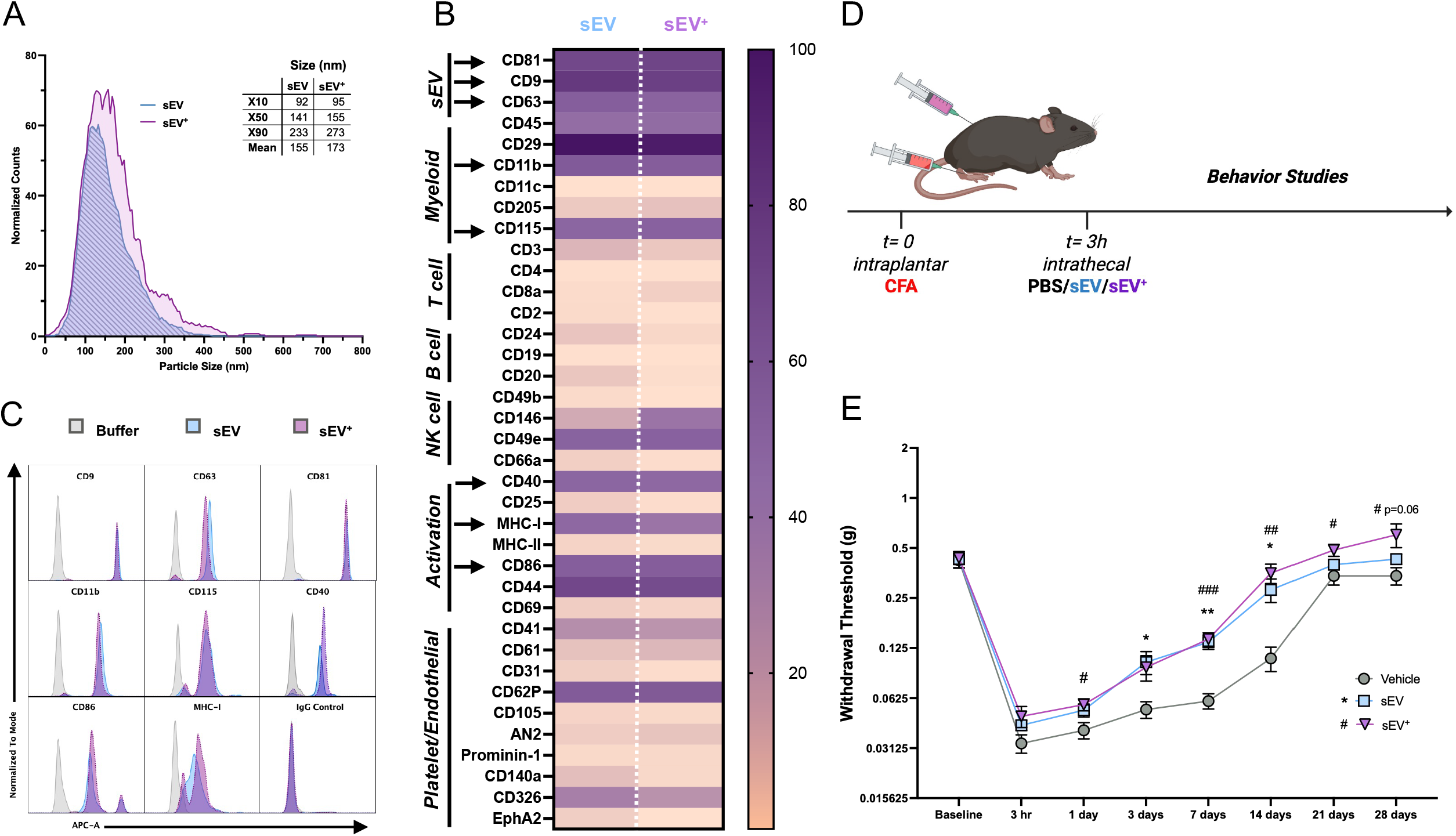
Macrophage derived small extracellular vesicles (sEVs) contain costimulatory molecules that could activate T cells and promote early resolution of inflammatory pain. sEVs were isolated from the media of RAW 264.7 macrophage cells cultured in complete media (DMEM + 10% sEV depleted FBS + 1% P/S) “sEV” or complete media with 1 µg/mL LPS “sEV^+^”. (**A**) Nanoparticle tracking analysis of sEVs showing the particle size distribution, consistent with the size of sEVs, including exosomes. (**B**) Weighted expression of surface immune markers on sEV populations using bead-based capture immuno-assay (MACSplex) with notable positive sEV, myeloid, and activation markers required to activate T cells (arrows) represented in (**C**) which shows the representative histograms of marker intensity as well as that of a negative IgG isotype control. (**D**) Schematic showing timing of intraplantar CFA, and intrathecal sEV administration (1 µg), followed by behavior testing. and (**E**) Mechanical allodynia evaluated using von Frey filaments applied to the plantar surface of the hind paw. Data shown as mean ± SEM (n=6-7), two-way repeated measures ANOVA with Dunnet’s test for multiple comparisons, * PBS vs. sEV, # PBS vs. sEV^+^. * *P*<0.05, ** *P*<0.01, *** *P*<0.001, sEV (derived from RAW 264.7 without LPS stimulation) shown in blue, and sEV^+^ (derived from RAW 264.7 with LPS stimulation) shown in purple.

For rigor, we reproduced our previous data, inducing inflammatory pain in male mice by injection of complete Freund’s adjuvant (CFA) into the hind paw and intrathecally administering either 1 µg sEV or sEV^+^ 3 hours after model induction as a therapeutic agent. We observed significant increased mechanical thresholds of sEV^+^ treated mice as measured by von Frey filaments, peaking at day 7 and 14 prior to model resolution, consistent with previous data **(Figure 1D-E, Supplemental Figure 1B)**. sEV showed similar results. We also tested this paradigm in female mice and observed similar results **(Supplemental Figure 1C-D)**. As the effects of sEV and sEV^+^ were similar in male and female, we chose to focus our studies on using male mice henceforth.

### sEV^+^ treatment of splenocyte cultures increases APC and T cell activation

We first asked if T cells were capable of being activated by sEVs with APCs present in co-culture. We incubated primary splenocytes with 1 µg sEV or sEV^+^ and examined the resulting activation of APCs and T cells after 24 hours. We confirmed that sEVs are capable of being taken up by T cells *in vitro* **(Supplemental Figure 2A-B)**. APCs of sEV^+^ treated splenocytes showed strong increases in activation including macrophages, B cells, and other myeloid populations (CD11b^+^F4/80^−^) as measured by upregulation of costimulatory molecules CD80 and CD86 on the surface **(Figure 2 A, C-D, Supplemental Figure 3 C-D)**. Additionally, both CD4^+^ and CD8^+^ T cells treated with sEV^+^ showed increased activation **(Figure 2 E-F, H-I, Supplemental Figure 3 G-H)**. We also measured if there were any changes in the percentage of Tc1, Th1 or Treg subtypes, and observed an increase in Treg (CD4^+^FOXP3^+^CD25^+^) cells(**Figure 2 G, J)** which was associated with increased T-bet expression **(Supplemental Figure 3 I)**. Non-macrophage myeloid cells (CD11b^+^F4/80^−^) decreased (**Figure 2B**) and total T cells increased **(Supplemental Figure 3 E-F)** within sEV^+^ treated splenocytes. In contrast to sEV^+^, sEV had minimal impact on APC or T cell activation. We also checked if T cell depletion affected APC activation (95% purity, **Supplemental Figure 4 J**) and observed similar upregulation of CD80 and CD86 across APC populations in T cell depleted splenocyte cultures **(Supplemental Figure 4 A-H)**.

**Figure 2.**
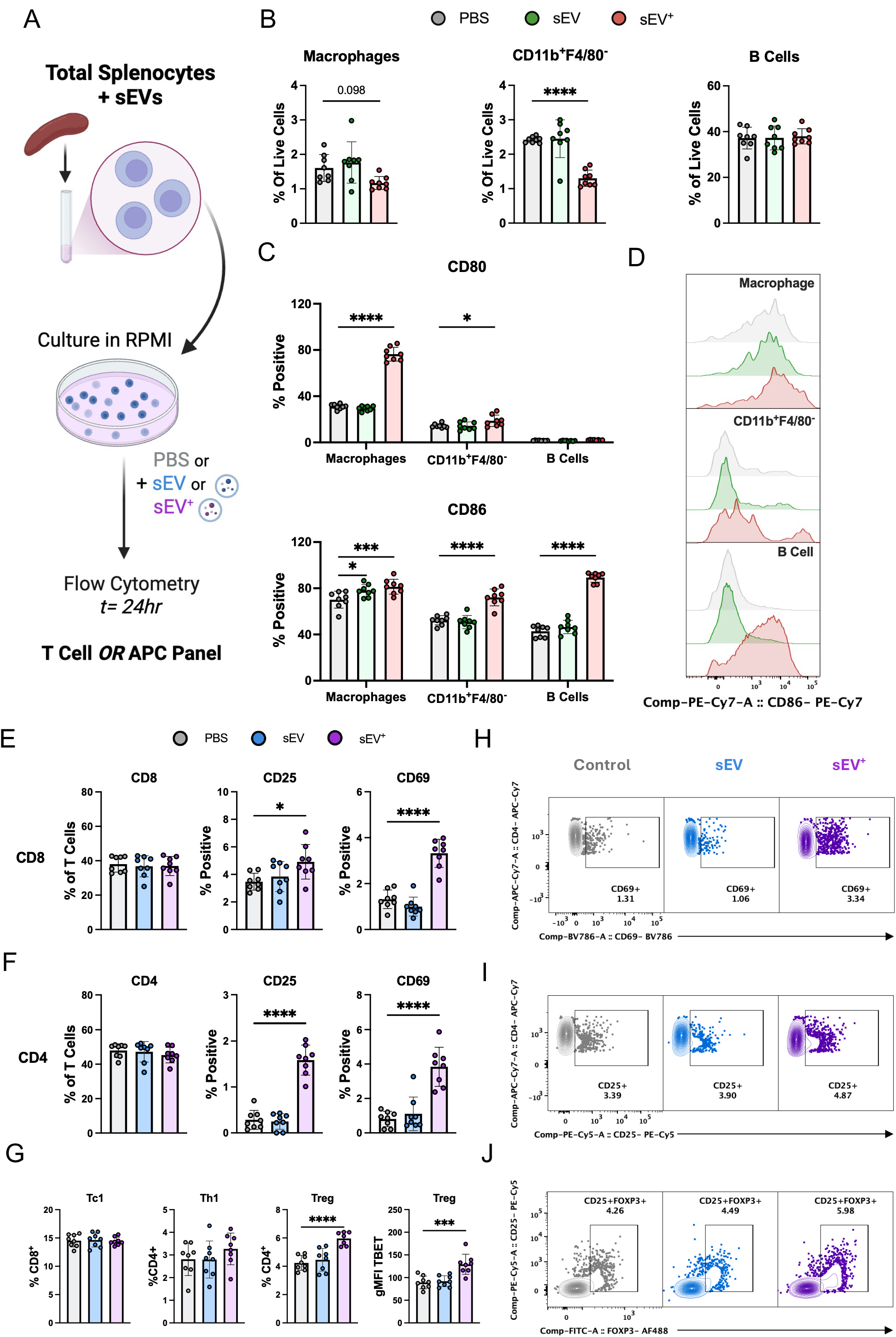
sEV^+^ treatment increases activation of antigen presenting cells (APCs) and T cells in splenocyte cultures. (**A**) Schematic for *in vitro* experiments using total splenocytes cultured for 24 hours with or without sEVs (1 µg) to determine indirect effects of sEVs on T cells. (**B**) Indirect effects of sEVs on APCs in splenocyte cultures. Effects of sEV treatment on APC percentages within the culture, including macrophages (CD11b^+^F4/80^+^), CD11b^+^F4/80^−^ (monocytes, dendritic cells, and neutrophils), and B Cells (CD11b^−^CD19^+^). One-way ANOVA with Dunnet’s test for multiple comparisons. **** *P*<0.0001. (**C**) Expression of co-stimulation and activation markers CD80 (top) and CD86 (bottom) on splenocyte APC populations. Two-way ANOVA with Dunnet’s test for multiple comparisons * P<0.05, *** *P*<0.001, **** *P*<0.0001. (**D**) Histogram of CD86 expression. PBS-grey, sEV-green, sEV^+^-red. (**E**) Indirect effects of sEVs on T cells in splenocyte cultures. Effects of sEV treatment on CD8^+^ and (**F**) CD4^+^ percentages within the culture as well as percentage of cells expressing early (CD69) and late (CD25) activation markers. (**G**) Effects of sEV treatment on T cell polarization in splenocyte cultures showing Tc1 (CD8^+^T-Bet^+^), Th1 (CD4^+^T-Bet^+^), and Treg (CD4^+^FOXP3^+^), as well as gMFI of T-Bet in Treg populations. Mean ± SD (n=8), one-way ANOVA with Dunnet’s test for multiple comparisons, single outlier removed by Grubbs with alpha= 0.05. * *P*<0.05, **** *P*<0.0001. (**H-I**) Concatenated bivariate flow contour plots of CD4^+^ from **F** showing increased CD69 (top) and CD25 (bottom) expression on sEV^+^ treated cells. (**J**) Concatenated bivariate flow contour plots of CD4^+^ T cells from **G** showing increased expression of Treg (CD25^+^FOXP3^+^) in sEV^+^ treated cells. PBS-grey, sEV-blue, sEV+-purple.

### Direct T cell activation by sEVs is modest compared to APC-dependent activation induced by sEV^**+**^

As sEVs activated T cells in splenocyte cultures, it was unclear if this occurred through direct interaction (independent of APCs) or indirectly (through APC-mediated uptake and presentation). Since sEVs contained the necessary surface markers to directly activate T cells, we cultured primary T cells (CD3^+^, 95% purity, **Supplemental Figure 4 J**) with sEV or sEV^+^ and examined expression of early (CD69) and late (CD25) activation markers at 4 and 24 hours post-sEV addition. sEVs activated T cells at low frequencies, with sEV significantly increasing activation in CD8^+^ T cells at 4 hours while sEV^+^ showed increased CD4^+^ activation at 24 hours **(Figure 3A-C)**. We also examined if sEVs altered T cell activation under concurrent exogenous stimulation by α-CD3/CD28. Both sEV and sEV^+^ showed significant increases in CD8^+^ T cell activation at 24 hours, while CD4^+^ activation was increased at both 4 and 24 hours. At 24 hours, only sEV^+^ treatment resulted in a significant decrease in the percentage of CD8^+^ T cells in culture **(Figure 3D-F)**. Additionally, only sEV^+^ treated T cells exhibited reduced MFI of CD69 and CD25 among positive cells, despite an increased proportion of activated cells **(Supplemental Figure 2A-D)**. These data demonstrate that while both sEV and sEV^+^ can directly activate T cells the effects are modest when not paired with concurrent α-CD3/CD28 stimulation. Instead, only sEV^+^ are capable of robust stimulation of APCs and indirect activation of T cells in splenocyte cultures.

**Figure 3.**
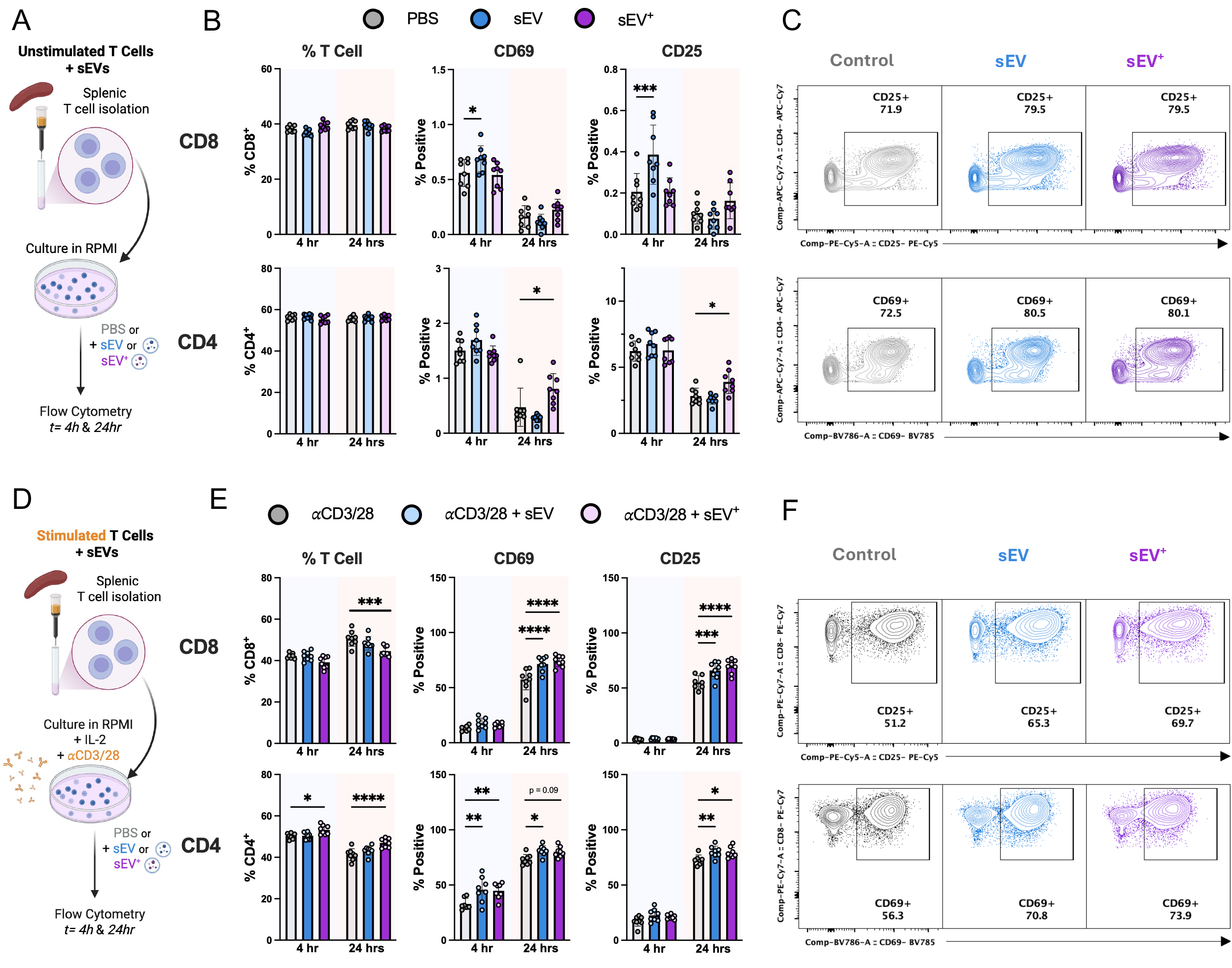
sEVs are capable of both directly activating and potentiating T cell activation. (**A**) Schematic for *in vitro* experiments using splenic T cells isolated by negative magnetic selection and cultured for 4 or 24 hours with or without sEVs (1 µg) to determine the direct effects of sEVs. (**B**) The effects of sEV treatment alone on CD8^+^ (top) and CD4^+^ (bottom) percentages within the culture as well as percentage of cells expressing early (CD69) and late (CD25) activation markers. Mean ± SD (n=8), two-way ANOVA with Dunnet’s test for multiple comparisons * *P*<0.05, *** *P*<0.001. (**C**) Concatenated bivariate flow contour plots of CD4^+^ cells at 24 hours from **B** showing increased CD25 (top) and CD69 (bottom) expression on sEV^+^ treated cells. (**D**) Schematic for *in vitro* experiments as in **A** but stimulating with α-CD3/28 and IL-2 to examine if sEVs potentiate T cell activation. (**E**) The effects of sEV treatment of stimulated T cells on CD8^+^ (top) and CD4^+^ (bottom) percentages within the culture as well as percentage of cells expressing early (CD69) and late (CD25) activation markers. Mean ± SD (n=8), two-way ANOVA with Dunnet’s test for multiple comparisons. * *P*<0.05, ** *P*<0.01, *** *P*<0.001, **** *P*<0.0001. (**F**) Concatenated bivariate flow contour plots of CD8^+^ cells at 24 hours from **E** showing increased CD25 (top) and CD69 (bottom) expression on sEV^+^ treated cells. sEV shown in blue, and sEV^+^ shown in purple.

### sEVs suppress Treg polarization and promote Th1 polarization both directly and indirectly

In splenocyte cultures sEV^+^ treatment increased Treg frequency, suggesting sEVs may alter the polarization and function of activated T cells. We examined whether sEVs contain TGF-β, a key signal for Treg polarization. ELISA and SMAD2 phosphorylation studies both confirmed the presence of TGF-β in sEVs **(Supplemental Figure 6A-K)**.

We next sought to determine if sEV treatment of T cells could directly alter polarization toward Treg and Th1 subtypes. Naïve CD4^+^ T cells (CD4^+^CD62L^+^CD44^−^CD25^−^, 95% purity, **Supplemental Figure 7C**) were stimulated with α-CD3/CD28 in the presence of IL-2 and treated with sEVs. The following day, increasing doses of TGF-β or IL-12 were added to the cultures. Polarization and IFN-γ secretion were measured on day 4. Surprisingly, sEV and sEV^+^ treated T cells showed decreased Treg polarization (FOXP3^+^) at low doses of TGF-β. In contrast, sEV and sEV^+^ treated T cells increased Th1 polarization (T-Bet^+^) at low to mid doses of IL-12, and at mid to high doses produced significantly more IFN*γ***(Figure 4A-E)**.

**Figure 4.**
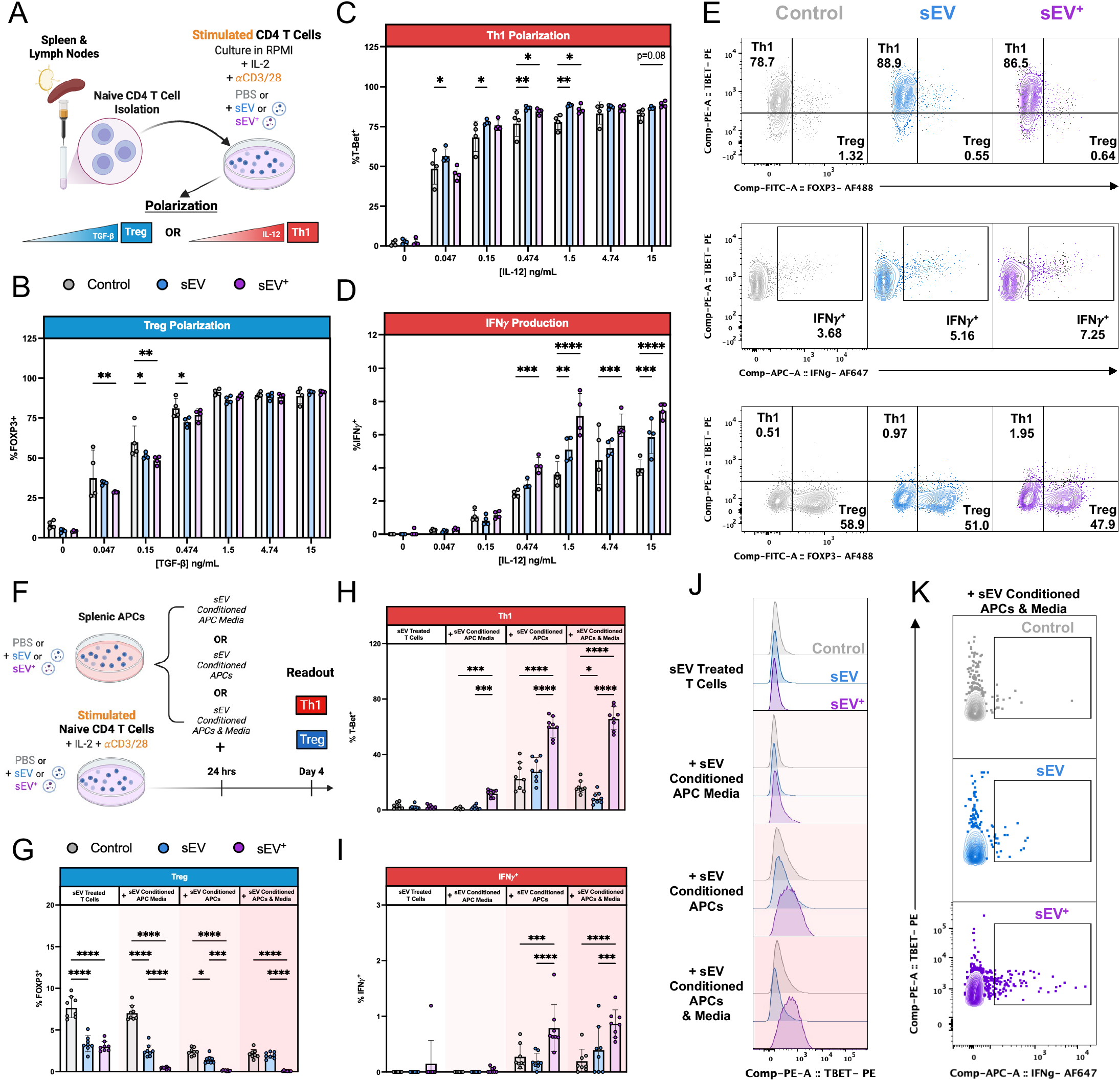
sEV^+^ suppresses Treg and promotes Th1 polarization through both direct and indirect effects. (**A**) Schematic for *in vitro* polarization experiments in which naïve CD4^+^ T cells of naïve mice were stimulated with −CD3/28, IL-2 and varying concentrations of TGF-β (Treg) or IL-12 (Th1) for 2 days with or without sEVs (1 µg), and removed from stimulation on day 2 and supplemented with IL-2 each day until day 4 when they were restimulated (α-CD3/28) and analyzed. (**B**) Quantification of Treg polarization (T-Bet^−^FOXP3^+^) in sEV treated cultures in the presence of increasing TGF-β concentrations. (**C**) Quantification of Th1 polarization (T-Bet^+^FOXP3^−^) in sEV treated cultures in the presence of increasing IL-12 concentrations. (**D**) Quantification of IFN*γ*production of Th1 cells from **C**. Mean ± SD (n=4), two-way ANOVA with Dunnet’s test for multiple comparisons * *P*<0.05, ** *P*<0.01, *** *P*<0.001, **** *P*<0.0001. (**E**) Concatenated bivariate flow contour plots of Th1 polarized cells treated with 1.5 ng/mL IL-12 showing increased T-Bet^+^ cells (top) and IFN*γ*(middle) production. Treg cultures treated with 0.150 ng/mL TGF-β (bottom) show decreased FOXP3^+^ cells. (**F**) Schematic for *in vitro* polarization experiments where the effects of sEV or sEV^+^ conditioned APCs or APC media on T cell polarization were examined. Conditioned media, APCs, or both were supplemented to T cell cultures at 24 hours in a matched fashion for APC/T cell cultures (PBS/PBS, sEV/sEV, sEV^+^/sEV^+^). (**G**) Quantification of Treg polarization (T-Bet^−^FOXP3^+^) in sEV treated cultures with or without conditioned APCs and media. (**H**) Quantification of Th1 polarization (T-Bet^+^FOXP3^−^) in sEV treated cultures with or without conditioned APCs and media and (**I**) IFN*γ*production of Th1 cells from **H**. Mean ± SD (n=8), two-way ANOVA with Dunnet’s test for multiple comparisons showing within group comparisons * *P*<0.05, ** *P*<0.01, *** *P*<0.001, **** *P*<0.0001. (**J**) Concatenated histograms showing increased expression of T-Bet in T cells treated with sEV^+^ conditioned APC media and APCs from **H**. (**K**) Concatenated bivariate flow contour plots showing increased IFN expression in in T cells treated with sEV^+^ conditioned APC media and APCs from **I**. sEV shown in blue, and sEV^+^ shown in purple.

Because this outcome differed from that observed in short-term splenocyte cultures **(Figure 4G)**, we next examined whether indirect, APC-mediated effects of sEVs influenced T cell polarization. We treated APCs (T cell–depleted splenocytes) in parallel with sEV or sEV^+^. After day 1, instead of adding TGF-β or IL-12, naïve T cells received conditioned media, APCs, or both from sEV-treated APC cultures. **(Figure 4F)**. We observed that while direct sEV treatment of T cells was sufficient to reduce Treg polarization at low levels, both soluble mediators and cell interactions of sEV^+^ treated APCs significantly reduced Treg polarization**(Figure 4G, Supplemental Figure 8B)**. In addition, only conditioned media from sEV^+^ treated APCs increased Th1 polarization and IFN*γ*production, with sEV^+^ treated APCs themselves producing a more pronounced effect **(Figure 4H-K, Supplemental Figure 8B)**. Taken together these results demonstrate that while both sEV and sEV^+^ directly suppress Treg and promote Th1 polarization, sEV^+^ treated APCs have a greater contribution to these changes through both soluble factors and cell–cell interactions.

### sEV^+^ suppress T cell and promote B cell proliferation

As sEV^+^ treated stimulated T cells showed alterations in CD4^+^/CD8^+^ percentages at 24 hours **(Figure 3E)** and increased total T cells in 24-hour splenocyte cultures **(Supplemental Figure 3F)**, we next tested whether sEVs altered T cell proliferation. CFSE labelled T cells were stimulated with α-CD3/CD28 and IL-2 to induce proliferation and treated with sEV/sEV^+^. Proliferation was evaluated 3 days after stimulation. Both CD4^+^ and CD8^+^ T cells in sEV and sEV^+^ treated cultures showed a significant reduction in proliferation as measured by the proliferation index and differences in generational percentages, while the effects of sEV^+^ were stronger **(Figure 5A-D)**. Although treatment of CFSE-labeled splenocyte cultures with sEV or sEV^+^ alone was insufficient to induce detectable T cell proliferation above background (data not shown), sEV^+^ treatment did increase the percentage of dividing B cells within the culture (**Figure 5E)**. This increase in B cell proliferation was also observed in cultures of CFSE labelled APCs (T cell depleted splenocytes) **(Figure 5F)**.

**Figure 5.**
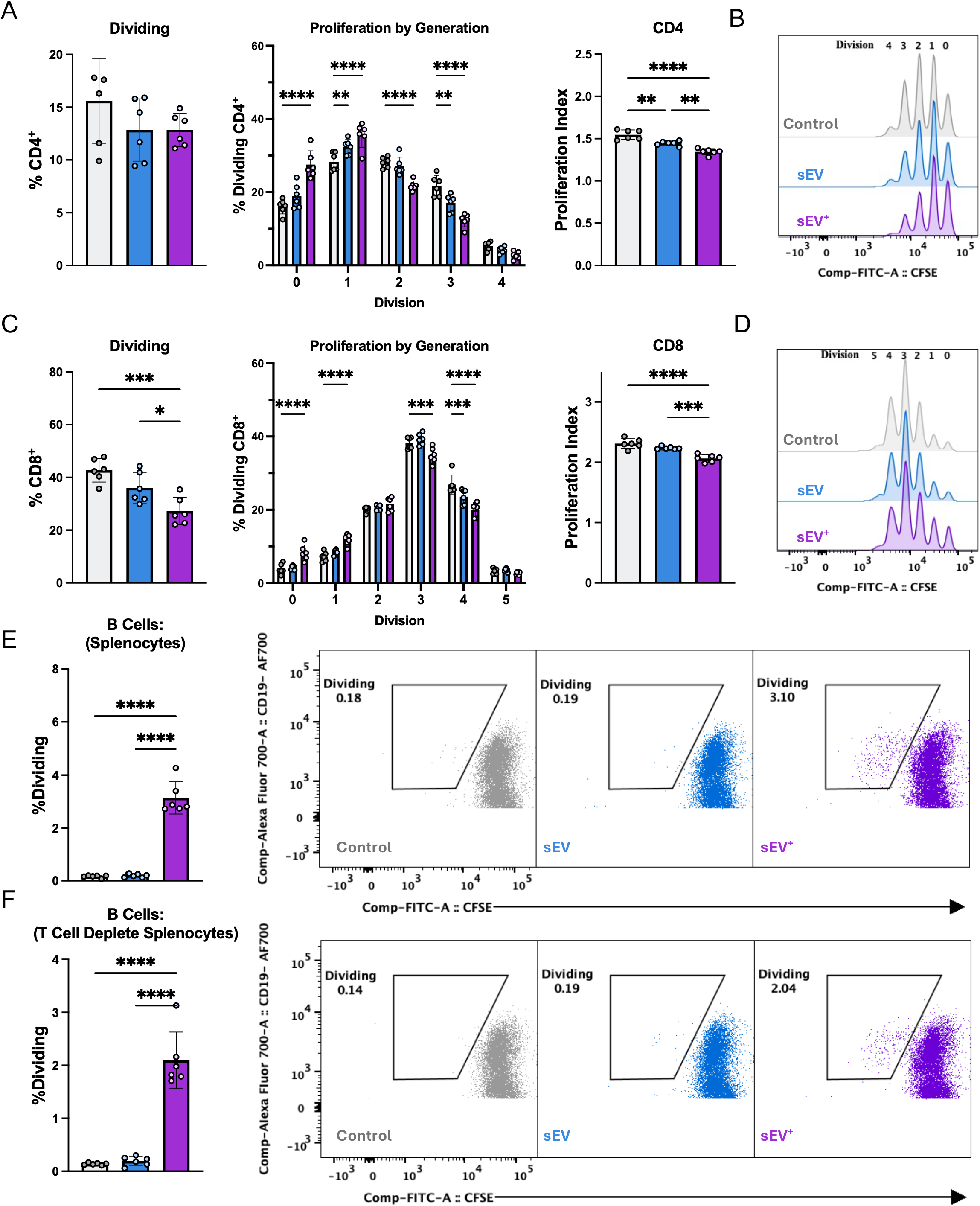
sEV^+^ suppresses T cell and promotes B cell proliferation. Proliferation of stimulated T cells (-CD3/28) that had been stained with CFSE to track daughter cell generations was measured 3 days after stimulation. (**A**) Proliferation of CD4^+^ T cells showing the total % dividing (left), % of dividing within each daughter generation/division (middle) and the proliferation index (right). (**B**) Concatenated histograms of the CFSE staining of CD4^+^ T cells with decreasing CFSE intensity in each successive daughter generation. (**C**) Proliferation of CD8^+^ T cells, showing the total % dividing (left), % of dividing within each daughter generation/division (middle) and the proliferation index (right). Mean ± SD (n=6), one or two-way ANOVA with Dunnet’s test for multiple comparisons * *P*<0.05, ** *P*<0.01, *** *P*<0.001, **** *P*<0.0001. (**D**) Concatenated histograms of the CFSE staining of CD8^+^ T cells, with decreasing CFSE intensity in each successive daughter generation. (**E**) Cell proliferation of CFSE labeled B cells in mixed splenocyte cultures and concatenated bivariate flow dot plots (right) showing increased CFSE low B cells. (**F**) T cell depleted splenocytes with percent dividing (left) and concatenated bivariate flow dot plots (right) showing increased CFSE low B cells. Mean ± SD (n=8), one-way ANOVA with Dunnet’s test for multiple comparisons, **** *P*<0.0001. sEV shown in blue, and sEV^+^ shown in purple.

### Late-phase resolution of CFA inflammatory pain by therapeutic sEV^+^ treatment requires T cells

Given the potential of sEV^+^ to activate T cells and modulate their polarization and proliferation *in vitro*, we asked whether T cells were required for the accelerated pain resolution mediated by therapeutic intrathecal sEV^+^ administration. We utilized *Rag2*^*−/−*^ mice, which lack both T and B cells. We adoptively transferred total CD3^+^ T cells from wild-type mice into *Rag2*^*−/−*^ recipients two weeks prior to CFA injury, enabling assessment of the requirement of T cells independent of B cell-dependent interactions **(Figure 6A)**. Adoptive transfer was confirmed at five weeks after completion of behavioral studies **(Figure 6B-C)**. We observed that at 3 days post CFA injection, both sEV^+^ treated *Rag2*^−/−^ mice and *Rag2*^*−*^*/*^*−*^ mice that received adoptively transferred T cells had significantly higher mechanical withdrawal thresholds than *Rag2*^*−/−*^ mice that had not received T cells. In contrast, only sEV^+^ treated *Rag2*^*−/−*^ that received adoptively transferred T cells showed significantly higher pain thresholds at 7 and 14 days post CFA **(Figure 6D, Supplemental Figure 9)**. Thus, T cells are required for sEV^+^ mediated late-phase inflammatory pain resolution, demonstrating the importance of T cells for sEV^+^-mediated effects observed at 7 days post-CFA and after.

**Figure 6.**
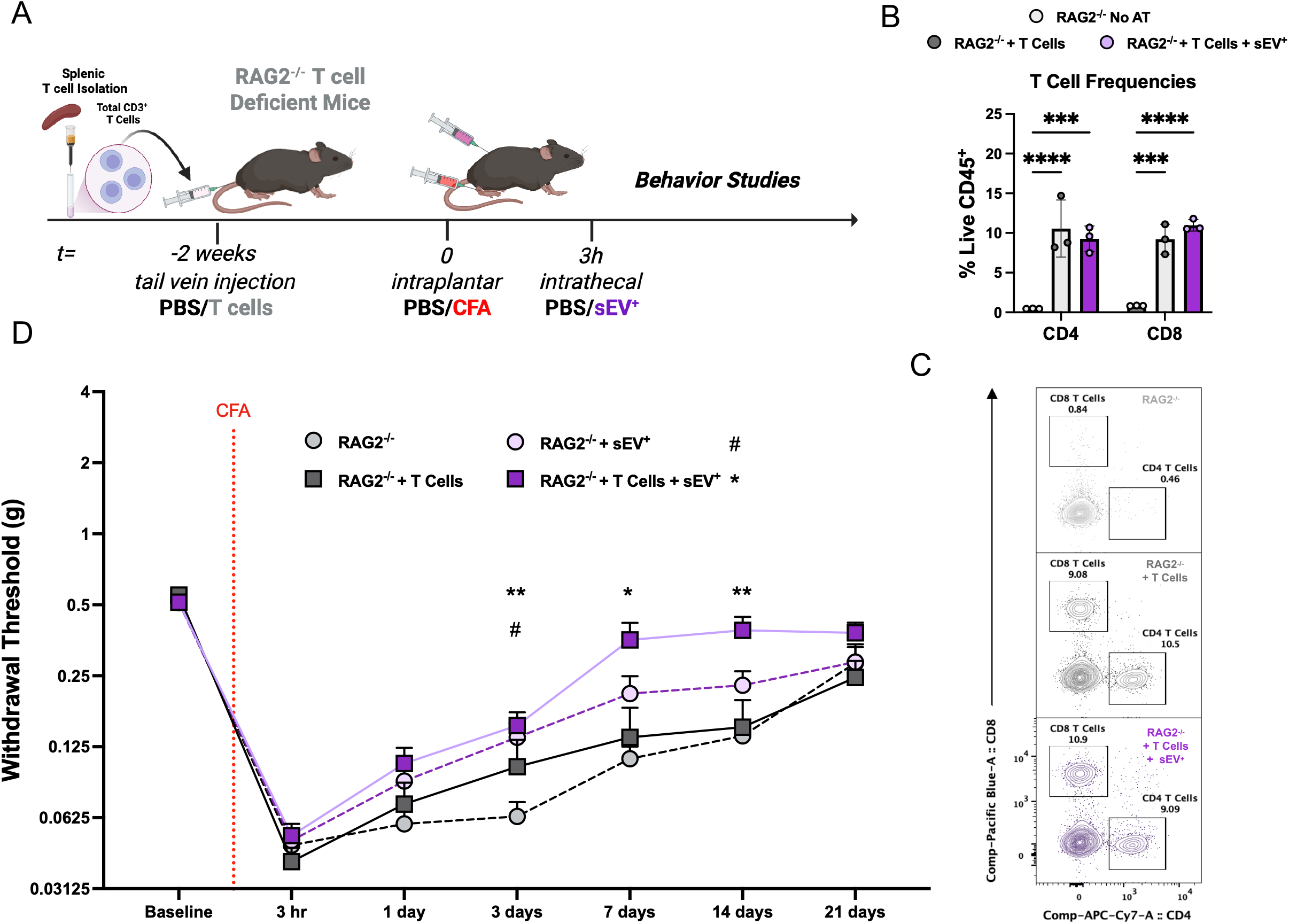
Late-phase inflammatory pain resolution mediated by intrathecal delivery of sEV^+^ requires T cells. (**A**) Schematic showing design for adoptive transfer of total CD3^+^ T cells isolated from WT C57BL/6J mice by negative magnetic selection of splenocytes and injected into the tail vein of T cell deficient *Rag2*^*–/–*^ mice. Two weeks after adoptive transfer, mice were subjected to *intraplantar* CFA administration and *intrathecal* sEV administration (1 µg) and assessed for mechanical sensitivity of the injected paw up to 3 weeks. (**B**) Confirmation of T cell adoptive transfer in splenocytes of mice 3 weeks after CFA to ensure T cell presence through the duration of the experiment, visualized as concatenated bivariate flow contour plots in (**C**). (**D**) von Frey test evaluating mechanical sensitivity of *Rag2*^−/−^ mice with or without T cell transfer and treatment with sEV^+^. Mean ± SEM (n=7-10), two-way repeated measures ANOVA with Dunnet’s test for multiple comparisons. * *P*<0.05, ** *P*<0.01 # *Rag2*^−/−^ vs *Rag2*^−/−^ + sEV^+^, * *Rag2*^−/−^ vs *Rag2*^−/−^ + T cells + sEV^+^.

### Therapeutic intrathecal delivery of sEV^+^ has minimal effects on splenic immune populations but strongly activates lymphocyte populations in the cervical draining lymph nodes

To evaluate how intrathecal delivery of sEV^+^ may modulate immune responses *in vivo*, we first assessed if systemic circulating immune responses were affected by sEV^+^ treatment. We evaluated splenic immune populations at early (1 day) and late (7 days) timepoints post-CFA and sEV^+^ administration **(Supplementary Figure 10A)**. Overall, there were minimal sEV^+^ dependent effects on lymphoid and myeloid cell populations in the spleen **(Supplementary Figure 10B-G)**. Overall, while there were small changes in immune populations in the spleen, such as the reduced percentage of CD4^+^ T cells **(Supplementary Figure 10B)**, we concluded that intrathecal administration of sEV^+^ has minimal effects on circulating splenic immune cells.

We next examined if sEV^+^ may affect immune populations in local draining lymph nodes, a more direct target for their immunostimulatory potential[28]. While the major site of lymphatic drainage from the hindpaw is the popliteal lymph node (pLN), it also drains more distally to the sacral and internal iliac lymph nodes (siLN). Lymphatic drainage of intrathecally administered agents has been shown to drain to multiple sites, including the siLN as well as the superior cervical lymph nodes (scLN)[29, 30]. We evaluated immune cell populations at D1 and D7 from the pLN, siLN, and scLN to observe changes within LNs that exclusively drain from the paw (pLN-*red*) and intrathecal injection (scLN-*purple*) as well as a shared site (siLN) **(Figure 7A)**. Strikingly, we observed significantly increased total cell counts within the scLN on D1 driven exclusively by sEV^+^ treatment. Significantly increased cell counts within the pLN on D7 were driven by CFA treatment as expected **(Figure 7B)**. CFA and not sEV^+^ treatment drove significant decreases in the percentage of CD4^+^ and CD8^+^ T cells and a significant increase in the percentage of B cells in the pLN on D1. In the scLN however, the same effect was observed but was induced by sEV^+^ treatment alone and not CFA **(Figure 7C)**. Interestingly, although very low in total number, only sEV^+^ treatment drove increases in neutrophils in the scLN on D1, among all LNs and timepoints **(Supplemental Figure 11C)**. In myeloid populations of the pLN, CFA induced significant increases primarily in Ly6C^Hi^ monocytes on D1 and macrophages, Ly6C^Lo^ and Ly6C^Hi^ monocytes on D7 (**Figure 7D**). In the scLN sEV^+^ treatment drove significant increases of Ly6C^Lo^ monocytes and decreases in dendritic cells on D1, both of which were non-significant at D7. Intriguingly, the siLN which is a shared draining lymph node for CFA and sEV^+^ treatment, showed significant reduction of macrophages and Ly6C^Lo^ monocytes in sEV^+^ treated animals (**Figure 7D**).

**Figure 7.**
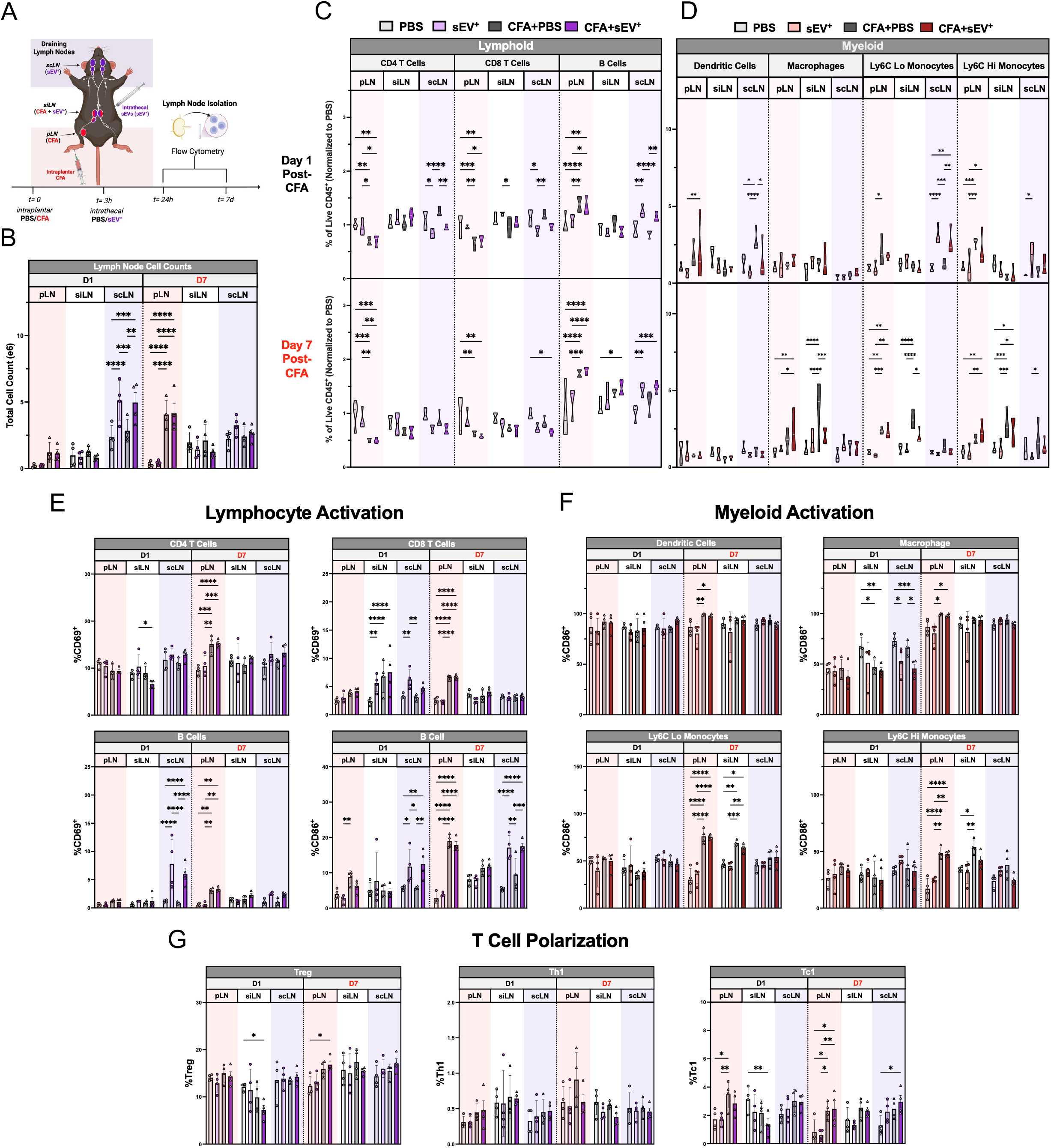
Intrathecal delivery of sEV^+^ strongly activate immune populations in the cervical draining lymph nodes. (**A**) Schematic showing design to evaluate differences in activation of immune cell populations in draining lymph nodes. Draining lymph nodes (LN) were assessed for intraplantar injection (popliteal (p)LN, sacral/internal iliac (si)LN) and intrathecal injection (superior cervical (sc)LN, and siLN) where pLN is unique to CFA injection (light red background), scLN is unique to sEV^+^ injection (light purple background) and the siLN represents an intersecting draining site (white background in figures). Immune cell populations from draining lymph nodes were assessed at 1 and 7 days after CFA/sEV^+^ injection to assess lymphoid (PBS-light grey, sEV^+^-light purple, CFA+PBS-dark grey, CFA+sEV^+^-dark purple) and myeloid (PBS-light grey, sEV^+^-light red, CFA+PBS-dark grey, CFA+sEV^+^-dark red) populations via flow cytometry. (**B**) Count of total cells in respective lymph nodes as measured by hemocytometer at D1 (left) and D7 (right). Truncated violin plots showing the percentage of lymphoid (**C**) and myeloid (**D**) cell populations in lymph nodes at day 1 (top) and day 7 (bottom) post CFA/sEV^+^ administration. Cell percentages were normalized by cell type to PBS control. Lymphoid and myeloid were separated due to differences in variance as lymphoid represent >95% of cells in the LN while myeloid is <5% (Supplemental Figure 11); (n=4) two-way ANOVA with Dunnet’s test for multiple comparisons * *P*<0.05, ** *P*<0.01, *** *P*<0.001, **** *P*<0.0001. (**E**) Lymphocyte activation as measured by expression of early activation marker CD69 across lymphocyte populations and additionally co-stimulation marker CD86 for B cells. (**F**) Myeloid activation as measured by expression of co-stimulatory signal CD86 across macrophages (CD11b^+^Ly6G^−^Ly6C^−^F4/80^+^), Ly6C^Lo^ and Ly6C^Hi^ monocytes (CD11b^+^Ly6G^−^Ly6C^Lo/Hi^), and dendritic cells (CD11b^+^Ly6G^−^Ly6C^−^ CD11c^+^MHCII^+^). (**G**) T cell polarization as measured by percentage of Treg (CD4^+^FOXP3^+^), Th1 (CD4^+^T-Bet^+^), and Tc1 (CD8^+^T-Bet^+^). (n=4) two-way ANOVA with Dunnet’s test for multiple comparisons. * *P*<0.05, ** *P*<0.01, *** *P*<0.001, **** *P*<0.0001.

The changes in lymphocyte numbers induced by sEV^+^ treatment in the scLN on D1 were accompanied by significantly increased activation of CD8^+^ T cells and B cells as measured by expression of CD69 (T & B cells) and CD86 (B Cells). By D7 only the increases in B cell activation persisted **(Figure 7E)**. In the scLN CD4^+^ T cells showed non-significant increases in Th1 and Treg on D1 and D7 respectively. Tc1 cells were highest in sEV^+^ treated CFA animals **(Figure 7G**). Increased activation induced in the pLN by CFA was seen at D7 across T and B cells. Of note is that sEV^+^ treatment alone activated CD8^+^ T cells in the siLN on D1 and showed the lowest CD4^+^ activation in sEV^+^ treated CFA mice **(Figure 7E)**. This was paralleled in T cell polarization data, where the siLN on D1 showed the lowest number of Tregs and Tc1 cells in sEV^+^ treated CFA mice **(Figure 7G)**. CFA treated mice had increased myeloid activation across all cell types on D7 in the pLN as measured by CD86 expression, matching the broad myeloid activation across cell types observed by their increased percentages **(Figure 7F)**. The only significant change in myeloid activation observed in the scLN was a reduction in macrophage activation at D1. We also observed that while CFA treatment alone increased Ly6C^Hi^ monocyte activation in the siLN on D7, sEV^+^ treated CFA animals did not. General trends in alternate activation markers of lymphocytes (CD25) and myeloid populations (MHCII) were observed as well, though to a much smaller degree **(Supplemental Figure 11D-E)**.

In summary, we observed that intraplantar CFA and intrathecal sEV^+^ were accompanied by broad increases and activation of lymphocyte and myeloid populations that were exclusive to their draining lymph nodes, the pLN and scLN respectively. Intriguingly, while myeloid cell populations were altered in the scLN by sEV^+^, this was not accompanied by increased expression of costimulatory marker CD86. In fact, CD86 decreased on macrophages, while sEV^+^ only increased T and B cell activation (CD69) and B cell expression of CD86. The siLN, the shared draining lymph node site, showed mixed effects of CFA and sEV^+^ treatment including increased CD8^+^ T cells, decreased Treg and Tc1 polarization subtypes on D1, as well as lower myeloid cell recruitment on D7.

## Discussion

Our findings show that macrophage-derived sEVs not only activate and modulate T cell responses, but that T cells are essential mediators of sEV-driven inflammatory pain resolution. We first replicated our earlier findings demonstrating that therapeutic intrathecal administration of sEVs and sEV^+^ effectively promoted the resolution of CFA-induced mechanical hypersensitivity[9]. The presence of APC-derived surface markers critical for T cell activation (MHC-I/II, CD86, CD40) within both sEV and sEV^+^ preparations supports their potential to directly engage the TCR and modulate T cell responses. While both sEV and sEV^+^ were able to directly activate T cells at low levels *in vitro*, in line with prior studies of dendritic cell–derived sEVs[7], the two vesicle populations differed in their ability to stimulate T cells. sEV were only capable of activating CD8^+^ T cells directly and transiently, while sEV^+^ primarily activated CD4^+^ T cells directly and at 24 hours. This did not align with MHC-I/II expression and likely reflects differences in peptide loading or other cargo. Additionally, while both sEV and sEV^+^ could potentiate T cell activation under exogenous stimulation, only sEV^+^ induced robust T cell activation and upregulation of APC co-stimulatory molecules in splenic cultures. This was seen in T cell polarization assays as well, where direct sEV or sEV^+^ treatment suppressed Treg and promoted Th1 polarization while only sEV^+^ showed robust promotion of Th1 polarization indirectly through APCs. While both sEV and sEV^+^ could promote early resolution of CFA induced inflammatory pain, sEV^+^ showed a significantly higher immunostimulatory potential than sEV overall. Though not significant, sEV^+^ showed faster resolution than sEVs and they may contain shared mechanisms greater than their differences. Based on their modest direct effects on T cells and pronounced APC-dependent immunostimulatory capacity, we focused *in vivo* studies on sEV^+^.

It is curious that the stronger immunostimulatory responses induced by sEV^+^, typically associated with pro-inflammatory responses, are beneficial in the context of accelerating pain resolution. While this may appear counterintuitive at first, it is in line with the current perspective that acute inflammation is essential to healing and regeneration, but is a matter of timing and location[16]. Recent studies have shown that individuals exhibiting signatures of acute neutrophil activation at the time of injury were less likely to experience chronic pain[31]. sEV^+^ was able to induce acute neutrophil recruitment in the scLN at D1, though systemic alterations were not apparent. Picco et al. demonstrated the ability of intrathecally administered macrophage-derived sEVs to attenuate pain in a neuropathic pain model. In this model however, sEVs necessitated engineered loading of an antagomir against pro-inflammatory miRNA-155 and were suggested to target sensory neuron populations in the dorsal root ganglion (DRG)[32]. They reported increased levels of miR-155, miR-146, and miR-21 in sEVs derived from LPS-polarized M1-like macrophages, consistent with our previous findings showing similar upregulation of miR-146 and miR-21 in sEV^+^[9, 33]. Together, these findings suggest initial inflammatory responses induced by sEV^+^ may act to initiate alterations in systemic or regional inflammatory responses that modulate nociceptive signaling potentially shaping the downstream immune response.

Although T cells are increasingly recognized as modulators of pain, their exact role remains unclear, with studies across injury models and pain conditions reporting variable outcomes. Several groups have observed T cells to be dispensable for the establishment of CFA-induced inflammatory pain and post-operative pain models[19, 34], while others have reported that T cell deficiency decreased baseline pain thresholds and increased sensitivities to chemically induced inflammatory pain[35]. The role of T cells in the establishment of neuropathic pain models has also been mixed[36–38]. There is, however, a strong convergence on the importance of T cells for mediating resolution of pain in several different models. Rescue experiments using adoptive transfer of CD3^+^ T cells into *Rag2*^−/−^ knockout mice[19] and CD4^+^ effector T cells into nude mice[22], demonstrated an important role for T cells in the resolution of CFA-induced inflammatory pain. While one study observed the necessity of T cells for the resolution of CFA-induced inflammatory pain[19], another found T cells promoted resolution but were not strictly necessary[22], our results align more with the latter. Intriguingly, that study further showed adoptive transfer of activated Th1 cells had anti-nociceptive effects, supporting a functional role for pro-inflammatory effector T cells in modulating inflammatory pain.

While our studies could not definitively observe alterations in Tregs and Th1 cells in sEV^+^ treated animals *in vivo*, we demonstrated that sEV^+^ suppress Treg and promote Th1 polarization *in vitro*. This was contrary to our initial expectations, as Treg frequency was increased in splenocyte cultures 24 hours after sEV^+^ addition, and we detected small but potent amounts of TGF-β within sEVs. Intriguingly, the Treg population expanded in sEV^+^ treated APC cultures showed increased T-Bet^+^ expression, suggesting potential functional heterogeneity consistent with type 1 Treg cells[39]. However, we did not detect significant changes in complementary *in vitro* assays or in our *in vivo* analyses, leaving the functional relevance of this subset unresolved. T cell polarization likely varies across anatomical sites over time as it is influenced by the location of initial antigen encounter, identity of the APC and strength of its interaction[40], as well as the local microenvironment in the tissues to which T cells subsequently migrate. Another group reported that Treg inhibit Th1 responses at the site of peripheral nerve injury to counteract neuropathic pain[24]. Future studies will need to more precisely define the timing, location, and trafficking of these different cell populations to better understand their functional contributions. Our *in vivo* studies observed the strongest effect of sEV^+^ on the increased activation of CD8^+^ T cells in the shared CFA and sEV^+^ draining sacral and internal iliac lymph nodes. This was accompanied by the lowest percentages of Treg and Tc1 on D1 in sEV^+^ treated CFA animals. Previous reports have identified that CD8^+^ T cells are necessary for the resolution of chemotherapy-induced neuropathic pain through the secretion of IL-13[25, 27]. Furthermore, they showed the education of CD8^+^ T cells by prior cisplatin exposure was necessary for mediating this resolution[26]. The importance of prior training or antigen exposure in supporting pain resolution was also demonstrated in CD4^+^ studies mentioned above[21, 22]. It is plausible that sEV^+^ may act *in vivo* to prime T cells, directing them toward pro-resolving functional programs. Previous work has implicated the production of IL-13, IL-10[25, 27], and enkephalins[20, 22] as key mediators of these pro-resolving processes.

It remains unclear whether the sEV^+^-dependent promotion of pain resolution requires antigen-specific T cell receptor (TCR) engagement or can occur through bystander T cell responses. The prominence of effector T cell responses suggests prior TCR engagement. Adoptive transfer of educated CD8^+^ T cells from wild-type or transgenic TCR OT-I mice, whose T cells predominantly express an ovalbumin (OVA)-specific TCR, did not reveal a significant difference in inflammatory pain resolution[26]. However, because OT-I mice retain functional *Rag* genes, a residual population of T cells expressing non-transgenic TCRs can still develop[41]. In addition, the OT-I TCR exhibits imperfect antigen discrimination[42] and inflammatory conditions are associated with increased presentation of self-peptides and neo-epitopes[43, 44], complicating interpretation of antigen specificity in this setting. Male mice receiving wild-type compared with OT-I–educated CD8^+^ T cells showed a non-significant trend toward faster pain resolution[26]. Accordingly, the relative contributions of antigen-specific versus antigen-independent T cell responses cannot be definitively resolved and warrants further investigation.

This raises the question of antigen specificity regarding sEV-induced T cell responses. Identifying specific antigens from the complex contents of sEVs will prove challenging. One candidate lies in examining alloreactivity of sEVs from different sources. RAW 264.7 sEVs are derived from BALB/C mice with the H-2^d^ haplotype, while C5Bl/6J mice have the H-2^b^ haplotype. Alloreactive T cells are activated by allogeneic sEVs *in vitro* and *in vivo*, often necessitating allogeneic APCs to mediate their activation through cross-dressing or uptake and processing of donor sEVs[45, 46]. We observed robust activation of APC responses *in vivo* and *in vitro*, and these greatly amplified the activation of T cells by sEV^+^. As both sEV and sEV^+^ promote inflammatory pain resolution, but only sEV^+^ induced robust effects in context of APCs *in vitro*, allogenicity is likely not the only factor that contributes to *in vivo* T cell responses. However, studies have found that allogeneic sEVs loaded with OVA peptide were more effective at inducing antigen-specific T cell responses against OVA expressing tumors[47]. Allogeneic cells have also been shown to preferentially promote Th1 polarization[48], a finding consistent with our *in vitro* observations. Collectively, these observations suggest that alloimmune activation during systemic inflammation may enhance T cell responses more broadly, thereby contributing to sEV-mediated inflammatory pain resolution.

Sex is an important biological variable in pain studies and has significant implications for underlying mechanisms and outcomes. One study identified T cells as being important mediators to neuropathic pain only in female mice[49]. Another group identified Treg as a critical mediator of pain induced by intrathecal injection of CSF1 in only female mice[23]. We employed a model of inflammatory, not neuropathic pain, and did not observe sex-dependent differences in sEV-mediated pain resolution; however, the study was not powered or designed to formally detect sex differences. Focusing on male mice allowed us to narrow our study, as the RAW 264.7 cell line is derived from male mice, avoiding complications of female reactivity to minor histocompatibility antigens such as the H-Y antigen reported previously[50]. Future studies examining antigen-specific T cell responses of sEVs should consider the sex of sEV donor and recipient to better understand how expression of minor histocompatibility antigens may affect T cell activation, inflammatory responses, and pain resolution.

Finally, we provide the first evidence that intrathecal administration of sEV^+^ elicits robust activation of lymphocytes and APCs in intrathecal-draining scLNs. Meningeal and CNS-draining lymphatic pathways have emerged as areas of intense interest, challenging the long-held view that the CNS is an immune-privileged site. The cervical, sacral, and internal iliac LNs have been identified as major clearance sites of cerebrospinal fluid (CSF), with multiple studies demonstrating accumulation of intrathecally administered tracers[30, 51–53] including recent evidence of bacterial extracellular vesicle accumulation[54]. We observed an sEV^+^-dependent increase in total cell numbers in the scLN at day 1 post-injection. sEV^+^ treatment decreased the proportion of T cells while increasing B cells, consistent with our *in vitro* findings showing reduced T cell proliferation and enhanced B cell expansion. Notably, sEV^+^ also reduced dendritic cell frequencies while increasing monocytes and neutrophils in the scLN, suggesting B cells and Ly6C^Lo^ monocytes may represent prominent APC populations targeted by sEV^+^. In contrast, sEV^+^ effects were not evident in the CFA-draining pLN, indicating spatial constraints of sEV^+^-effects in the lymphatics. Indeed, we were able to observe mixed effects of sEV^+^ and CFA in the shared draining siLNs, including alterations in lymphocyte proportions, T cell activation, and shifts in macrophage and Ly6C^Hi^ monocyte populations. It is worth noting that the results from our *Rag2*^−/−^ studies utilize mice that lack mature B cells, suggesting there may be functional redundancy in APC populations and the effect of sEV^+^ on monocytes may be critical for future studies. Collectively, these findings suggest sEV^+^ can modulate complex systemic inflammatory and immune responses at regionally shared sites.

Deficiencies of lymphatic clearance of CSF increase with age[52] and case reports describing improvements in chronic pain following manual promotion of lymphatic drainage at sites distant from the original injury[55, 56] suggest an important role for lymphatic function in chronic pain conditions. Notably, one group demonstrated that T cells originating from distant lymphatic sites are required to migrate to the leptomeninges of lumbar dorsal roots to drive the transition from acute to chronic pain following tibial nerve injury[38]. The migration of T cells from the scLN following activation is of considerable interest for future therapeutic strategies. Effector and memory T cells are known to exit LN and re-enter systemic circulation[28, 57]. Although we observed minimal changes in splenic immune populations, our study may not have been sufficiently powered to detect modest systemic effects. Notably, sEV^+^ treatment of CFA mice did not increase total splenic CD4^+^ T cell numbers, indicating systemic immune modulation may occur through altered trafficking rather than numerical expansion. Supporting this idea, a recent study demonstrated that localized LN responses to topical antigens can influence distant allergic responses to orally ingested food antigens through immune cell trafficking and circulating secreted factors[58]. Future studies may also consider neuronal modulation as the cervical and meningeal lymphatic network has been shown to permit retrograde transport of nanoparticles to the brain and along peripheral nerves[59], in agreement with the observations in the DRG[32]. Consistent with this, neuronal regulation of antigen flow within lymphatics has been reported[60] as has descending modulation of peripheral immune responses through vagal projections in neuropathic pain[61]. Together, these findings highlight multiple potential mechanisms by which sEV^+^ may exert its effects.

While T cells are established mediators of pain resolution, strategies to therapeutically harness these endogenous mechanisms without relying on cell-based interventions remain limited. Our findings suggest that sEVs may represent a novel, cell-free approach to engage endogenous T cell-mediated pain resolving pathways without the cost, complexity, or logistical barriers associated with personalized cellular therapies. Furthermore, the capacity of sEVs to modulate immune responses within cervical LNs further suggests broader immunomodulatory potential, with implications extending beyond pain to cancer and inflammatory, and autoimmune diseases.

## Materials and Methods

### Cell culture and sEV isolation

RAW 264.7 mouse macrophage cells (ATCC TIB-71) were cultured in complete media (1x Dulbecco’s Modified Eagle Medium (DMEM)) (Sigma-Aldrich) supplemented with 10% heat-inactivated Fetal Bovine Serum (FBS) (Corning) and 100 U/mL penicillin-streptomycin (Pen-Strep) (Thermofisher) at 37°C with 5% CO_2_. At 60-70% confluence, RAW 264.7 cells were washed with 1X Phosphate Buffered Saline (PBS) depleted of ions; complete media was changed to complete media made with exosome-deplete FBS, in the presence (sEV^+^) or absence (sEV) of 1 μg/mL Lipopolysaccharide (LPS) (Sigma-Aldrich). At 24 hours conditioned media was collected and centrifuged at 500 ×g for 10 minutes at 4°C to pellet dead cells, then 12,000 ×g for 30 minutes at 4°C to remove debris and large vesicles. The supernatant was filtered through a 0.22 μm syringe filter and 12 mL of filtered supernatant was concentrated to a final volume of 500 µL with 100 kDa concentrator (Amicon, Millipore Sigma) by centrifugation at 5000 ×g at 4°C for 30 minutes in a fixed angle rotor (Beckman Coulter). A 35 nm qEV size exclusion chromatography column (Izon ICO-35) was washed using 1X PBS and 500 µL of concentrated supernatant was then loaded onto the column followed by 2.5 mL PBS to collect 3 mL void volume. When the column flow stopped, 2 mL of PBS was added to collect EV zone sample. The EV volume (2 mL) was centrifuged for 70 minutes at 110,000 ×g at 4°C. The EV pellet was resuspended in DPBS, assessed for protein concentration using microBCA, and stored at −80°C until use.

### Nanoparticle tracking analysis (NTA)

Size distribution and concentration of sEVs was assessed with a NanoSight LM10 instrument, equipped with a 405 nm laser, that tracks Brownian motion under laser illumination (Malvern Panalytical). Samples diluted in 0.1 µm filtered PBS were injected into the sample chamber and further diluted to a concentration of 20-90 particles/frame. Five 60-second videos were captured for each sample.

### Characterization of sEV surface markers

The mouse MACSPlex exosome kit (Miltenyi Biotec) was used for detection of EV specific markers per manufacturer’s instructions. Briefly, sEVs were diluted to 120 μL in MACSPlex Buffer. The sEVs were incubated with 15 μL of MACSPlex Exosome capture beads overnight at room temperature using an orbital shaker (450 rpm). The following day, samples were washed with 500 μL of MACSPlex Buffer and centrifuged at 3000 ×g for 5 minutes. After aspirating 500 μL of the supernatant, 15 μL of MACSPlex Exosome detection reagent (CD9, CD63, or CD81 cocktail) was added to each tube and incubated for an hour using an orbital shaker. After washing and centrifugation, the tubes were incubated for 15 minutes at room temperature on an orbital shaker. MACSPlex exosome capture beads with Evs were resuspended by pipetting up and down, and analyzed by flow cytometry for various markers, including isotype control. Positive populations were gated and final %Pos was calculated by subtracting a buffer control sample. For representation of relative marker intensities, samples were weighted and displayed as %Pos if <50%, and if ≥50% were expressed as (%Pos*0.5)+((%Pos*0.5)*(MFI/MFI of highest sample).

### Animal studies

Male or female C57Bl/6J (WT) and *Rag2*^−/−^ mice without T or B cells (8–12 weeks old) (Jackson laboratory, #008449) were maintained in a 12-hour light/dark cycle and provided food and water *ad libitum. Rag2*^−/−^ mice were housed in the barrier facility. Behavior tests were performed in accordance with the NIH guidelines for the Care and Use of Laboratory Animals and approved by the Institutional Animal Care & Use Committee of Drexel University College of Medicine. For sEV treatment, mice received an intrathecal injection of 1 µg of sEVs three hours after CFA administration. Using a Hamilton syringe and 30-gauge needle, 10 µl of sEVs or PBS was injected under the dura mater at the lumbar enlargement.

### CFA model of inflammatory pain

Persistent peripheral inflammation was induced by administration of 50% emulsified complete Freund’s adjuvant (CFA) (Sigma) in PBS into the plantar surface of right hind paw of mice (20 μl). Mechanical paw thresholds were determined by applying a series of ascending force von Frey filaments (Bioseb) to the hind paw[9]. An average of three baseline measurements were made prior to CFA injection. All behavior studies were performed by researchers blinded to the treatment received.

### Isolation of primary splenocytes, T cells, and APCs

Spleens were collected and processed into single cell suspensions by passing spleens through a 70 µm cell strainer. Cells were centrifuged at 350 ×g for 5 minutes at 4°C. Cell pellet was resuspended in 3 mL of 1x RBC lysis buffer (eBioscience) for one minute at room temperature. RBC lysis was inactivated by adding 10 mL of complete media (RPMI 1640 + 1% P/S + 10% FBS). Cells were centrifuged at 350 ×g for 5 minutes at 4°C. Resulting splenocytes were resuspended in either complete media for experiments or isolation buffer (1x PBS + 0.5 % BSA + 2 mM EDTA) for downstream processing via magnetic isolation into Total CD3^+^ T cells or APCs (T cell depleted fractions) using the pan T cell Isolation Kit (Miltenyi Biotec, 130-095-130) according to manufacturer protocol. Briefly, splenocytes labelled with a cocktail of biotin-conjugated monoclonal antibodies (CD11b, CD11c, CD19, CD45R (B220), CD49b (DX5), CD105, MHC-II, and Ter-119) were bound to magnetic anti-biotin micro beads and passed through a LS column. Unlabeled T cells were collected in the resulting flow-through while the magnetically labeled fraction (T cell depleted) was retained in the magnetic column and subsequently removed from the magnet and eluted. Naïve CD4^+^ T cells were isolated using a similar procedure as above but pooling lymph nodes and spleen at the start and using the naïve CD4^+^ T cell isolation kit (Miltenyi Biotec, 130-104-453).

### Culture of primary total T cells

For primary T cell cultures, 1×10^5^ total T cells were cultured in 0.1 mL of complete T cell media (RPMI 1640 + 1 mM HEPES + 1 mM sodium pyruvate + 5 µM βME + 1% P/S + 10% FBS) in 96-well plates (NEST, #701011) for 4 or 24 hours. For stimulated T cell cultures, plates were pre-coated with −CD3 and −CD28 (Biolegend) diluted in PBS (100 µL at 1 µg/mL each) for 3-4 hours at 37℃ and washed with 100 µL PBS once prior to plating. Stimulated cultures were supplemented with 30 U/mL IL-2 (R&D Systems, #202-IL). To assess the effect of sEVs, cultures were treated with 1 µg of sEV or sEV^+^. T cell activation was assessed by flow cytometry (**Supplemental Table 1**).

### Culture of primary splenocytes and APCs

Primary splenocytes or APCs (T cell depleted splenocytes) were cultured in 96-well plates in complete media (RPMI 1640 + 1% P/S + 10% FBS) for 24 hours (1×10^5^ cells in 0.1 mL). To assess the effect of sEVs, cultures were treated with 1 µg of sEV or sEV^+^. T cell and APC activation was assessed by flow cytometry (**Supplemental Table 1**).

### Labeling of sEVs and uptake studies

sEVs or cells were labeled with general lipophilic fluorescent membrane dyes, PKH67 and PKH26 respectively, according to manufacturer’s instructions (Sigma-Aldrich). Briefly, 20 µg of sEVs were diluted in 1 mL of diluent buffer. An equivalent volume of PBS was diluted in 1 mL of diluent buffer to serve as dye control. Diluted PKH67 dye was added to sEVs or PBS and incubated for 5 minutes in the dark at room temperature. Staining was stopped by addition of 1% BSA. Labeled sEVs were centrifuged for 70 minutes at 110,000 ×g at 4°C. The supernatant was discarded, and the pellet was resuspended in 2 mL of PBS and centrifuged again. sEV pellet was resuspended in 50 μL PBS and quantified using microBCA kit (Thermofisher) for protein estimation. Total T cells or APCs were washed twice with serum free RPMI 1640 media, centrifuging at 400 ×g for 5 minutes. Cells were stained with PKH26 as above with two washes in complete media at 400 ×g for 5 minutes. Cells were resuspended in RPMI 1640 media prepared with exosome-deplete FBS for treatment with sEVs. 18 mm coverslips (Fisherbrand) were placed on a 12 well plate (Corning) and 1×10^6^ PKH26 labeled (CD3^+^) T cells or APCs were treated with 1 µg PKH67 labeled sEVs per well or an equivalent amount of PBS dye control at 37°C for 4 hours. Cells were washed with 0.1 M PB twice and fixed with 4% PFA for 10 minutes at room temperature. The coverslips containing cells were mounted on glass slides using prolong gold antifade reagent (Life technologies) and dried overnight in the dark. The prepared glass slides were stored at 4°C. Cells were visualized with laser scanning confocal microscope (Confocal FV3000) and images were acquired using FluoView Imaging software (Olympus FV1000).

### Enzyme-linked immunosorbent assay (ELISA)

To determine cytokine concentrations, recipient cell culture supernatants were analyzed with a TGF-β1 ELISA kit (R&D Cat DY1679-05) according to manufacturer’s protocol.

### Assessment of sEV induced pSMAD2 in total T cells

To assess phosphorylation of pSMAD2, 4×10^5^ total T cells (CD3^+^) were plated in a 96-well deep well plate in 0.2 mL of serum free media (SFM, RPMI + 1% P/S + 1 mM HEPES + 1 mM Sodium Pyruvate + 5 µM βME). SFM was used as FBS contains TGF-β and affects basal levels of pSMAD2. Cells were left overnight and then treated with TGF-β or sEVs at various doses. For inhibitor experiments cells were pre-incubated with 10 µM ALK inhibitor (Selleck Chem, #SB-43154) or 20 µg/mL α-TGF-β (BioXcell, #BE0057) 1 hour prior to TGF-β or sEV addition. 1 hour after TGF-β or sEV addition, live dead stain was added to cells and they were spun down at 300 ×g for 5 minutes, media was aspirated, and cells were resuspended in 100 µL of ice-cold methanol and fixed at 4℃ for 10 min. Samples were washed and stained for pSMAD2 in flow buffer. This was then repeated for secondary antibody staining. Cells were then washed and resuspended for acquisition.

### CD4 T cell polarization assays

To assess the direct effect of sEVs on CD4 T cell polarization, 5×10^4^ naïve CD4 T cells were plated in 0.1 mL of complete T cell media (RPMI 1640 + 1 mM HEPES + 1 mM sodium pyruvate + 5 µM βME + 1% P/S + 10% FBS + 30 U/mL IL-2) in α-CD3/ α-CD28 coated 96-well plates. Cells were treated with or without 1 µg sEV or sEV^+^ for 24 hours. Cells were then treated with varying doses of IL-12 (Biolegend, #577002) or TGF-β (R&D Systems, #240B) to induce Th1 or Treg polarization. 24 hours later cells were replated off α-CD3/ α-CD28 stimulation and supplemented with IL-2. On the 4^th^ day cells were re-stimulated with soluble α-CD3/ α-CD28 (2 µg/mL) for 6 hours in the presence of golgi-inhibitor brefeldin (Biolegend) and assessed for polarization and IFN*γ*production by flow cytometry (**Supplemental Table 1**). To assess indirect effects of sEV treated APCs on T cell polarization, T cells were plated as above but in 75 µL, while the T cell deplete fractions were plated in parallel in 96-well plates in complete media (RPMI 1640 + 1% P/S + 10% FBS) for 24 hours (1×10^5^ cells in 0.1 mL). T cells and APCs were treated with 1 µg sEV, sEV^+^, or PBS. At 24 hours, APC cultures of each treatment were pooled (i.e. all sEV^+^) and centrifuged at 350 ×g for 5 min at 4℃. Supernatant (conditioned media) was removed and centrifuged again to pellet any remaining cells. T cells were treated with either 75 µL of conditioned media, or 5×10^4^ conditioned in APCs in 75 µL of fresh or conditioned media. T cells were only treated with conditioned APCs or media of like treatments (ie. sEV^+^ T cells → sEV^+^ APCs). 24 hours later cells were replated off α-CD3/ α-CD28 stimulation and supplemented with IL-2. On the 4^th^ day cells were supplemented with brefeldin for 6 hours and assessed for polarization and IFN*γ*production by flow cytometry (**Supplemental Table 1**).

### Flow Cytometry

Cells were transferred to 96-well deep well V-bottom plates and centrifuged at 350 ×g at 4°C for 5 minutes to pellet cells. Media was flicked off and cells were stained with live/dead stain in PBS for 15 minutes at 4°C. Cells washed with flow buffer (1x PBS + 2% FBS + 2 mM EDTA + 0.04% sodium azide) and centrifuged 350 ×g at 4°C for 5 minutes. Cells were blocked with Trustain FcX (Biolegend) for 5 minutes and then stained for any fixation sensitive markers (CD69/CD19) in flow buffer for 15 minutes at RT. Cells were washed with flow buffer and centrifuged 350 ×g at 4°C for 5 minutes. Cells were then fixed in IC fixation buffer (ThermoFisher) for 15 minutes and washed twice with flow buffer and centrifuged 650 ×g at 4°C for 7 minutes. Cells were stained with remaining antibodies in flow buffer overnight, washed with flow buffer and centrifuged 650 ×g at 4°C for 7 minutes and resuspended in flow buffer for acquisition. For intracellular staining (T-Bet/FOXP3/IFN*γ*) after initial fixation, cells were incubated in permeabilization buffer (Thermofisher) with Trustain FcX for 5 minutes, remaining antibodies diluted in permeabilization buffer were added to stain targets overnight. Cells were washed once with permeabilization buffer and once with flow buffer and resuspended in flow buffer for acquisition. Panels and antibodies can be found in (**Supplemental Table 1**), and gating strategies can be found in **Supplemental Figures 3I-J, 4I, 5E, and 10C**.

#### CFSE proliferation assay

T cells, APCs, or splenocytes were washed and resuspended in staining media (RPMI 1640 + 1% FBS + 1% P/S) at 2×10^7^ cells/mL. Cells were stained with an equal volume of a 10 µM CFSE solution, prepared from a 5 mM stock solution of Cell Trace CFSE (Molecular probes) diluted in serum free media such that final concentration of FBS was 0.5%. Cells were stained for 15 minutes at 37℃ and quenched with 3 volumes complete RPMI. Cells were pelleted by centrifugation at 350 ×g for 5 minutes and washed twice more in 10 mL of complete media. Cells were plated at 5×10^4^ cells per well in 96 well plates. T cells were stimulated as in prior assays and proliferation was measured on day 3 via flow cytometry (**Supplemental Table 1**). FlowJo 10.10.1 was used to model CFSE divisions and generation gates. *Adoptive transfer of T lymphocytes in Rag2*^*−/−*^ *mice*. Two weeks prior to administration of CFA and intrathecal delivery of sEVs, 8×10^6^ total T cells isolated from the spleens of age and sex matched WT mice were intravenously (IV) injected into the tail vein in 100 μL. Control mice received an IV injection of vehicle (PBS + 0.1% BSA). Grafting of adoptively transferred T cells was assessed by flow cytometry.

### Sex as a Biological Variable

Other than initial behavior findings that demonstrate sEVs accelerate pain resolution in females as well, the rest of our study was performed with male mice, including all *in vitro* experiments. It is unknown whether the rest of the findings are observed in female mice.

### Statistical Analysis

For *in vitro* and *in vivo* experiments, statistical analyses were performed using GraphPad Prism (GraphPad Software, version 10.6.1). Six to eight biological replicates were used for *in vitro* studies. Four mice/group were used for *in vivo* molecular studies and six to ten mice/group for behavior studies. All data are presented as averages ± SEM or SD and identified in figure legends. Data were analyzed using Student’s t test (for experiments with two groups), one-way ANOVA followed by Dunnett’s multiple comparison test (for experiments with three or more groups and one variable), two-way ANOVA followed by Dunnett’s post hoc test (for experiments with two or more groups with multiple variables tested), or repeated measures two-way ANOVA followed by Dunnett’s post hoc test for behavior studies. Differences between averages were considered statistically significant when *P*<0.05.

### Study Approval

All procedures were consistent with the NIH’s Guide for the Care and Use of Laboratory Animals and the Ethical Issues of the International Association for the Study of Pain and were approved by the IACUC of Drexel University.

## Acknowledgments

We thank Dr. James M. Burns Jr. for helpful discussions and critical reading of the manuscript.

## Supplemental Table

**Supplemental Table 1. Flow cytometry panel staining**. Table describing the specific antibodies and fluorophores utilized for each panel, the staining conditions, and dilutions used.

## Supplemental Figure Legends

**Supplemental Figure 1. Intrathecal delivery of macrophage derived sEVs promotes early resolution of inflammatory pain and in male and female WT mice**. (**A**) Schematic showing timing of intraplantar CFA administration and intrathecal sEV administration (1 µg) as well as timepoints to assess mechanical sensitivity of the injected paw. (**B**) von Frey data from wild type male C57BL/6J mice in Figure 1E showing individual data points. Mean ± SEM (n=6-7), two-way repeated measures ANOVA with Dunnet’s test for multiple comparisons, * *P*<0.05, ** *P* <0.01, *** p<0.001. sEV (derived from RAW 264.7 without LPS stimulation) shown in blue, and sEV^+^ (derived from RAW 264.7 with LPS stimulation). Paw withdrawal threshold data from wild type female C57BL/6J mice shown as summary line graph (**C**) or as individual data points (**D**). Mean ± SEM (n=6), two-way repeated measures ANOVA with Dunnet’s test for multiple comparisons. **C**) * PBS vs. sEV, # PBS vs. sEV^+^. * *P* <0.05, ** *P* <0.01, **** *P* <0.0001. sEV shown in blue, and sEV^+^ shown in purple.

**Supplemental Figure 2. sEVs are taken up by both T cells and antigen presenting cells (APCs) in vitro**. Total T cells or T cell depleted (APCs) were labeled with lipophilic dye PKH26 and cultured with either PKH67 labeled sEVs (1 µg) or a dye control (remaining supernatant after washing during sEV labeling to account for residual dye and aggregates) for four hours at 37℃, fixed, and washed. Confocal images showing uptake of PKH67 labeled sEVs in both T cells (**A**) and APCs (**B**). Magnification 100x, scale 16 µm.

**Supplemental Figure 3. sEV**^**+**^ **treatment alone increases APC and T cell activation in splenocyte cultures**. Indirect effects on APCs in splenocyte cultures (PBS-grey, sEV-green, sEV^+^-red). **(A & B**) Percentages of macrophages (CD11b^+^F4/80^+^) and CD11b^+^F4/80^−^ of CD11b^+^ cells. Mean ± SD (n=8), one-way ANOVA with Dunnet’s test for multiple comparisons * *P* <0.05, ** *P* <0.01, **** *P* <0.0001. (**C & D**) gMFI of positive populations in Figure 2C for CD80 and CD86 across APC populations. Two-way ANOVA with Dunnet’s test for multiple comparisons * *P* <0.05, *** *P* <0.001, **** *P* <0.0001. Indirect effects on T cells in splenocyte cultures 24 hours after treatment with 1 µg sEVs (PBS-grey, sEV-blue, sEV^+^-purple). (**E & F**) Percentage of live cells and total T cells (CD3^+^) in culture. (**G & H**) MFI of positive populations in Figure 2E & F of early (CD69) and late (CD25) activation markers. (**I**) Histogram of T-Bet expression in Treg from Figure 2G. Gating strategy for evaluation of T cells (**J**) and APC populations (**K**) in splenocyte cultures from Figure 2 and Supplemental Figure 3.

**Supplemental Figure 4. sEV**^**+**^ **shows similar effects on APCs when treating T cell depleted cultures**. Direct effects on APCs in T cell depleted cultures 24 hours after treatment with 1 µg sEVs (PBS-grey, sEV-green, sEV^+^-red). (**A**) Schematic for *in vitro* experiments using T cell depleted splenocytes cultured for 24 hours with or without sEVs (1 µg) to determine direct effects on APCS. (**B**) The effects of sEV treatment on APC percentages within the culture, including macrophages (CD11b^+^F4/80^+^), CD11b^+^F4/80^−^ (monocytes, dendritic cells, and 41 neutrophils), and B cells (CD11b^−^CD19^+^). One-way ANOVA with Dunnet’s test for multiple comparisons * p<0.05. (**C**) Expression of co-stimulation and activation markers CD80 (Top) and CD86 (Bottom) on APC populations in culture. Two-way ANOVA with Dunnet’s test for multiple comparisons ** *P*<0.01, *** *P* <0.001, **** *P* <0.0001. (**D**) Concatenated histogram plots of APC populations from **C** showing increased CD86 expression on sEV^+^ treated cells. (**E & F**) Percentages of macrophages (CD11b^+^F4/80^+^) and CD11b^+^F4/80^−^ of CD11b^+^ cells. Mean ± SD *(n=8)*, one-way ANOVA with Dunnet’s test for multiple comparisons * *P* <0.05, ** *P* <0.01, **** *P* <0.0001. (**G & H**) gMFI of positive populations in D for CD80 and CD86 across APC populations. Two-way ANOVA with Dunnet’s test for multiple comparisons * *P* <0.05, **** *P* <0.0001. (**I**) Gating strategy for evaluation of APC populations in T cell deplete cultures. (**J**) Purity of MACS isolation of total T cell isolation showing input, negative fraction (total T cell enriched, CD3^+^TCRβ^+^) and positive fraction (APC Enriched).

**Supplemental Figure 5. sEVs are capable of both directly activating and potentiating T cell activation**. (**A**) Schematic for *in vitro* experiments using splenic T cells isolated by negative magnetic selection and cultured for 4 or 24 hours with or without sEVs (1 µg) to determine direct effects. (**B**) The effects of sEV treatment alone on CD8^+^ (Top) and CD4 (Bottom) expression *of* early (CD69) and late (CD25) activation markers as measured by gMFI of positive cell populations from Figure 3B. Mean ± SD (n=8), two-way ANOVA with Dunnet’s test for multiple comparisons. (**C**) Schematic for *in vitro* experiments as in A but stimulated with α-CD3/28 and IL-2 to examine if sEVs potentiate T cell activation. (**D**) The effects of sEV treatment of stimulated T cells on CD8^+^ (Top) and CD4^+^ (Bottom) expression *of* early (CD69) and late (CD25) activation markers as measured by gMFI of positive cell populations from Figure 3E. Mean ± SD (n=8), two-way ANOVA with Dunnet’s test for multiple comparisons **** *P*<0.0001. (**E**) Gating strategy for experiments in Figure 3 and Supplemental Figure 5 with FMOS for terminal markers shown in red. sEV shown in blue, and sEV^+^ shown in purple.

**Supplemental Figure 6. Macrophage derived sEVs contain TGF-β and induce pSMAD2 signaling**. (**A**) Amount of total and active TGF-β per µg of sEVs as measured by ELISA. Two-way ANOVA with Dunnet’s test for multiple comparisons (n=4) * *P* <0.05, **** *P* <0.0001. (**B**) Schematic for experiments measuring pSMAD2 levels in T cells. Total T cells were isolated from splenocytes of naïve male mice via negative magnetic depletion and subsequently cultured in serum free media (as TGF-β exists at high concentrations in serum) for 16 hours over night. The subsequent day, in inhibitor experiments, cells were pre-treated with α-TGF-β or ALK inhibitor for one-hour prior to the addition of sEVs or TGF-β as a positive control. At 1-hour post-treatment cells were immediately washed and methanol-fixed and stained for pSMAD2 signaling induced by TGF-β signaling in CD4^+^ and CD8^+^ cells by flow cytometry. (**C & E**) Representative dose response curves pSMAD2 signaling to TGF-β of in CD4^+^ and CD8^+^ T cells respectively, along with quantification of pSMAD2 MFI of sEV and sEV^+^ treated cells (**D**) and (**F**) at different doses. 4PL-curve fits along with one-way ANOVA with Dunnet’s test for multiple comparisons (n=3-6) * *P* <0.05, ** *P* <0.01, *** *P* <0.001, **** *P* <0.0001. Specificity of signal was tested by preincubation with either ALK inhibitor or α-TGF-β prior to TGF-β treatment with respective curves in CD4^+^ (**G**) and CD8^+^ (**I**) T cells. (**H & J**) show specificity of sEV signal at a dose of 3.16 µg sEVs4PL-curve fits along with one-way ANOVA with Dunnet’s test for multiple comparisons (n=3-6) * *P* <0.05, ** *P* <0.01, *** *P* <0.001, **** *P* <0.0001.

**Supplemental Figure 7. Gating strategies for T cell polarization experiments**. (**A**) Gating strategy and representative plots of CD4^+^ naïve T cells treated with high and low concentrations of IL-12 and TGF-β from Figure 4A-E. (**B**) Gating strategy and representative plots of CD4^+^ naïve T cells treated conditioned APCs and media Figure 4F-K. (**C**) Purity of MACS isolation of CD4 Naïve T cells showing input, negative fraction (Naïve CD4 enriched, CD4^+^TCRβ^+^CD62L^+^CD44^−^CD25^−^), and positive fraction (remainder).

**Supplemental Figure 8. sEV**^**+**^ **suppresses Treg and promotes Th1 polarization through both direct and indirect effects**. (**A**) Schematic for *in vitro* polarization experiments in which naïve CD4^+^ T cells isolated by negative magnetic selection from lymph nodes and spleens of naïve mice were stimulated with α-CD3/28 and IL-2, with or without sEVs for 24 hours. Additionally, APCs (T cell deplete) were also treated with or without sEVs for 24 hours. After 24 hours APC cultures were spun down and separated into conditioned media and APCs. Conditioned media, APCs, or both were supplemented to T cell cultures at 24 hours in a matched fashion for APC/T cell cultures (PBS/PBS, sEV/sEV, sEV^+^/sEV^+^). Cells were removed from stimulation on day 2 and supplemented with IL-2 each day until day 4 when they were restimulated (α-CD3/28) and analyzed. (**B**) Comparisons within treatment condition and between groups from **Figure 4G-I**. Mean ± SD (n=8), two-way ANOVA with Dunnet’s test for multiple comparisons * *P* <0.05, ** *P* <0.01, *** *P* <0.001, **** *P* <0.0001.

**Supplemental Figure 9. Late-phase inflammatory pain resolution mediated by intrathecal delivery of sEV**^+^ **requires T cells**. (**A**) Schematic showing design for adoptive transfer of total CD3^+^ T cells isolated from WT C57BL/6J mice by negative magnetic selection of splenocytes and injected into the tail vein of T cell deficient *Rag2*^−/−^ mice. Two weeks after adoptive transfer, mice were subjected to intraplantar CFA and intrathecal sEV administration (1 µg) and assessed for mechanical sensitivity of the injected paw via von Frey filaments for up to 3 weeks. (**B**) von Frey data evaluating mechanical sensitivity of *Rag2*^−/−^ mice with or without T cell transfer and treatment with sEV^+^ from Figure 6D showing individual data points. Mean ± SEM (n=7-10), two-way repeated measures ANOVA with Dunnet’s test for multiple comparisons * *P* <0.05, ** *P* <0.01.

**Supplemental Figure 10. Intrathecal delivery of sEV**^**+**^ **has minimal effects on splenic immune populations**. (**A**) Schematic showing design to evaluate splenic populations at 1 and 7 days after CFA/sEV^+^ administration via flow cytometry. (**B**) Truncated violin plots showing the percentage of lymphoid (left) (PBS-light grey, sEV^+^-light purple, CFA+PBS-dark grey, CFA+sEV^+^-dark purple) and myeloid (right) (PBS-light grey, sEV^+^-light red, CFA+PBS-dark grey, CFA+sEV^+^-dark red) cell populations in the spleen at day 1 (top) and day 7 (bottom) post CFA/sEV^+^ administration. Cell percentages were normalized by cell type to PBS control. Lymphoid and myeloid were separated due to differences in variance as lymphoid represent >90% of cells in the spleen while myeloid is <10% (Supplemental Figure 11B); (n=4) two-way ANOVA with Dunnet’s test for multiple comparisons. * *P*<0.05, ** *P*<0.01, *** *P*<0.001. (**C**) Lymphocyte activation as measured by expression of early activation marker CD69 across lymphocyte populations. (**D**) T cell polarization as measured by percentage of Treg (CD4^+^FOXP3^+^), Th1 (CD4^+^T-Bet^+^), and Tc1 (CD8^+^T-Bet^+^). (**E**) Myeloid activation as measured by expression of co-stimulatory signal CD86 across macrophages (CD11b^+^Ly6G^−^Ly6C^−^F4/80^+^), Ly6C^Lo^ and Ly6C^Hi^ monocytes (CD11b^+^Ly6G^−^ Ly6C^Lo/Hi^), and dendritic cells (CD11b^+^Ly6G^−^Ly6C^−^CD11c^+^MHCII^+^). Alternate activation markers for splenic lymphocyte (PBS-light grey, sEV^+^-light purple, CFA+PBS-dark grey, CFA+sEV^+^-dark purple) and myeloid (PBS-light grey, sEV^+^-light red, CFA+PBS-dark grey, CFA+sEV^+^-dark red) from A-E. (**F**) Lymphocyte activation as measured by expression of late activation marker CD25 (T cell) and alternate co-stimulatory molecule CD86 (B cell) across lymphocyte populations. (**G**) Myeloid activation as measured by expression of co-stimulatory signal CD86 across macrophages (CD11b^+^Ly6G^−^Ly6C^−^F4/80^+^) and Ly6C^Lo^ and Ly6C^Hi^ monocytes (CD11b^+^Ly6G^−^ Ly6C^Lo/Hi^). (**H**) Gating strategy for experiments from Supplemental Figure 10 & Figure 8 showing representative populations from the spleen with representative FMOs in red for terminal marker expression.

**Supplemental Figure 11. Intrathecal delivery of sEV**^**+**^ **strongly activates immune populations in the cervical draining lymph nodes**. (**A**) Schematic showing design to evaluate differences in immune cell activation of immune cell populations in draining lymph nodes. Draining lymph nodes (LN) were assessed for intraplantar injection (popliteal (p)LN, sacral/internal iliac (si)LN) and intrathecal injection (superior cervical (sc)LN, and siLN) where pLN is unique to CFA injection (light red background), scLN is unique to sEV^+^ injection (light purple background) and the siLN represents a potential intersecting draining site (overlap or white background in figures). Immune cell populations from draining lymph nodes were assessed at 1 and 7 days after CFA/sEV^+^ injection to assess lymphoid (PBS-light grey, sEV^+^-light purple, CFA+PBS-dark grey, CFA+sEV^+^-dark purple) and myeloid (PBS-light grey, sEV^+^-light red, CFA+PBS-dark grey, CFA+sEV^+^-dark red) populations via flow cytometry. (**B**) Parts of whole graphs showing the distribution of different cell populations across spleen and a representative lymph node (popliteal), where each circle represents 1%. (**C**) Neutrophils measured in lymph nodes at day 1 (Left) and day 7 (Right), which were analyzed separately due to the low % typically seen in LNs (<0.1%). (**D**) Alternate lymphocyte activation marker CD25 across T cell populations from Figure 7E. (**E**) Alternate myeloid activation and antigen presentation marker MHCII across macrophages (CD11b^+^Ly6G^−^Ly6C^−^F4/80^+^), and Ly6C^Lo^ and Ly6C^Hi^ monocytes (CD11b^+^Ly6G^−^ Ly6C^Lo/Hi^) from Figure 7F. (n=4) two-way ANOVA with Dunnet’s test for multiple comparisons * *P* <0.05, ** *P* <0.01, *** *P* <0.001, **** *P* <0.0001.

