## Supplementary material for "Macrophage-derived extracellular vesicles promote T cell–dependent inflammatory pain resolution": Supp. Table 1 and Supp. Figures 1-11

### Supplemental Table 1

| **T Cell Activation Panel** | | | | | | |
|---|---|---|---|---|---|---|
| **Target** | **Fluorophore** | **Clone** | **Vendor** | **Catalog Number** | **Step** | **Dilution** |
| **Live Dead Blue** | N/A | N/A | Thermo | L23105 | Live/Dead | 1000 |
| **CD8** | PE-Cy7 | QA17A07 | Biolegend | 155018 | Surface O/N | 1000 |
| **CD4** | APC-Cy7 | GK1.5 | Biolegend | 100414 | Surface O/N | 1000 |
| **CD25** | PE-Cy5 | PC61 | Biolegend | 102010 | Surface O/N | 1000 |
| **CD69** | BV785 | H1.2F3 | Biolegend | 104543 | Surface | 100 |
| **APC Activation Panel** | | | | | | |
| **Marker** | **Fluorophore** | **Clone** | **Vendor** | **Catalog Number** | **Step** | **Dilution** |
| **Live Dead Blue** | N/A | N/A | Thermo | L23105 | Live/Dead | 1000 |
| **CD3** | BV421 | 17A2 | Biolegend | 100228 | Surface O/N | 1000 |
| **CD19** | AF700 | eBioD3(1D3) | Thermofisher | 56-0193-82 | Surface | 33 |
| **CD11b** | APC | M1/70 | Biolegend | 101212 | Surface O/N | 1000 |
| **F4/80** | BUV737 | 1D3 | BD Bio | 612781 | Surface O/N | 1000 |
| **CD8** | PE-Cy7 | QA17A07 | Biolegend | 155018 | Surface O/N | 1000 |
| **CD4** | APC-Cy7 | GK1.5 | Biolegend | 100414 | Surface O/N | 1000 |
| **CD4 Polarization Panel** | | | | | | |
| **Marker** | **Fluorophore** | **Clone** | **Vendor** | **Catalog Number** | **Step** | **Dilution** |
| **Live Dead Blue** | N/A | N/A | Thermo | L23105 | Live/Dead | 1000 |
| **CD4** | BV785 | GK1.5 | Biolegend | 100453 | O/N post-Fix/Perm | 1000 |
| **TBET** | PE | 4B10 | Biolegend | 644810 | O/N post-Fix/Perm | 500 |
| **FOXP3** | AF488 | FJK-16S | Thermofisher | 53-5773-82 | O/N post-Fix/Perm | 500 |
| **IFNγ** | AF647 | XMG1.2 | BD Bioscience | 557735 | O/N post-Fix/Perm | 1000 |
| **pSMAD2 Studies** | | | | | | |
| **Marker** | **Fluorophore** | **Clone** | **Vendor** | **Catalog Number** | **Step** | **Dilution** |
| **Live Dead Zombie Yellow** | N/A | N/A | Biolegend | 423103 | Live/Dead | 300 |
| **CD4** | BV750 | GK1.5 | Biolegend | Biolegend | O/N post-Fix/Perm | 1000 |
| **CD8** | BV421 | QA17A07 | Biolegend | Biolegend | O/N post-Fix/Perm | 1000 |
| **pSMAD2** | Unconjugated (Rabbit) | SD207-1 | Thermofisher | 644810 | O/N post-Fix/Perm | 0.2 ug |
| **Isotype Control** | Unconjugated (Rabbit) | Polyclonal | Thermofisher | 02-6102 | O/N post-Fix/Perm | 0.2 ug |
| **Anti-Rabbit IgG** | PE | Polyclonal | Thermofisher | A10542 | O/N post-Fix/Perm | 1000 |
| **CFSE Studies** | | | | | | |
| **Marker** | **Fluorophore** | **Clone** | **Vendor** | **Catalog Number** | **Step** | **Dilution** |
| **Live Dead Blue** | N/A | N/A | Thermo | L23105 | Live/Dead | 1000 |
| **CFSE** | CFSE | N/A | Thermo | C34554 | Cell Labeling | 5 μM |
| **CD4** | BV785 | GK1.5 | Biolegend | 100453 | O/N post-Fix/Perm | 1000 |
| **TBET** | PE | 4B10 | Biolegend | 644810 | O/N post-Fix/Perm | 500 |
| **FOXP3** | AF488 | FJK-16S | Thermofisher | 53-5773-82 | O/N post-Fix/Perm | 500 |
| **IFNγ** | AF647 | XMG1.2 | BD Bioscience | 557735 | O/N post-Fix/Perm | 1000 |
| ***in vivo* Splencyte and Lymph Node Panel** | | | | | | |
| **Marker** | **Fluorophore** | **Clone** | **Vendor** | **Catalog Number** | **Step** | **Dilution** |
| **Live Dead Aqua** | N/A | N/A | Thermo | L34957 | Live/Dead | 1000 |
| **CD45** | BUV395 | 30-F11 | BD Bio | 564279 | Surface O/N | 500 |
| **CD11b** | APC | M1/70 | Biolegend | 101212 | Surface O/N | 500 |
| **Ly6G** | PercP-Cy5.5 | 1A8 | Biolegend | 127616 | Surface O/N | 5000 |
| **CD19** | AF700 | eBioD3(1D3) | Thermofisher | 56-0193-82 | Surface | 33 |
| **TCRβ** | BV421 | H57-597 | Biolegend | 109229 | Surface O/N | 200 |
| **CD8** | PE-Cy7 | QA17A07 | Biolegend | 155018 | Surface O/N | 500 |
| **CD4** | APC-Cy7 | GK1.5 | Biolegend | 100414 | Surface O/N | 500 |
| **CD25** | PE-Cy5 | PC61 | Biolegend | 102010 | Surface O/N | 1000 |
| **CD69** | BV785 | H1.2F3 | Biolegend | 104543 | Surface | 33 |
| **FOXP3** | AF488 | 4B10 | Biolegend | 644810 | O/N post-Fix/Perm | 500 |
| **TBET** | PE | FJK-16S | Thermofisher | 53-5773-82 | O/N post-Fix/Perm | 500 |
| **CD86** | BV650 | GL1 | BD Bio | 564200 | Surface O/N | 500 |
| **MHCII** | BV711 | M5/114.15.2 | Biolegend | 107643 | Surface O/N | 5000 |
| **Ly6C** | PE-Dazzle-594 | HK1.4 | Biolegend | 128044 | Surface O/N | 1000 |
| **CD11c** | RY703 | N418 | BD Bio | 759736 | Surface O/N | 500 |
| **F4/80** | BUV737 | 1D3 | BD Bio | 612781 | Surface O/N | 500 |

**Supplemental Table 1. Flow cytometry panel staining.** Table describing the specific antibodies and fluorophores utilized for each panel, the staining conditions, and dilutions used.

### Supplemental Figure 1

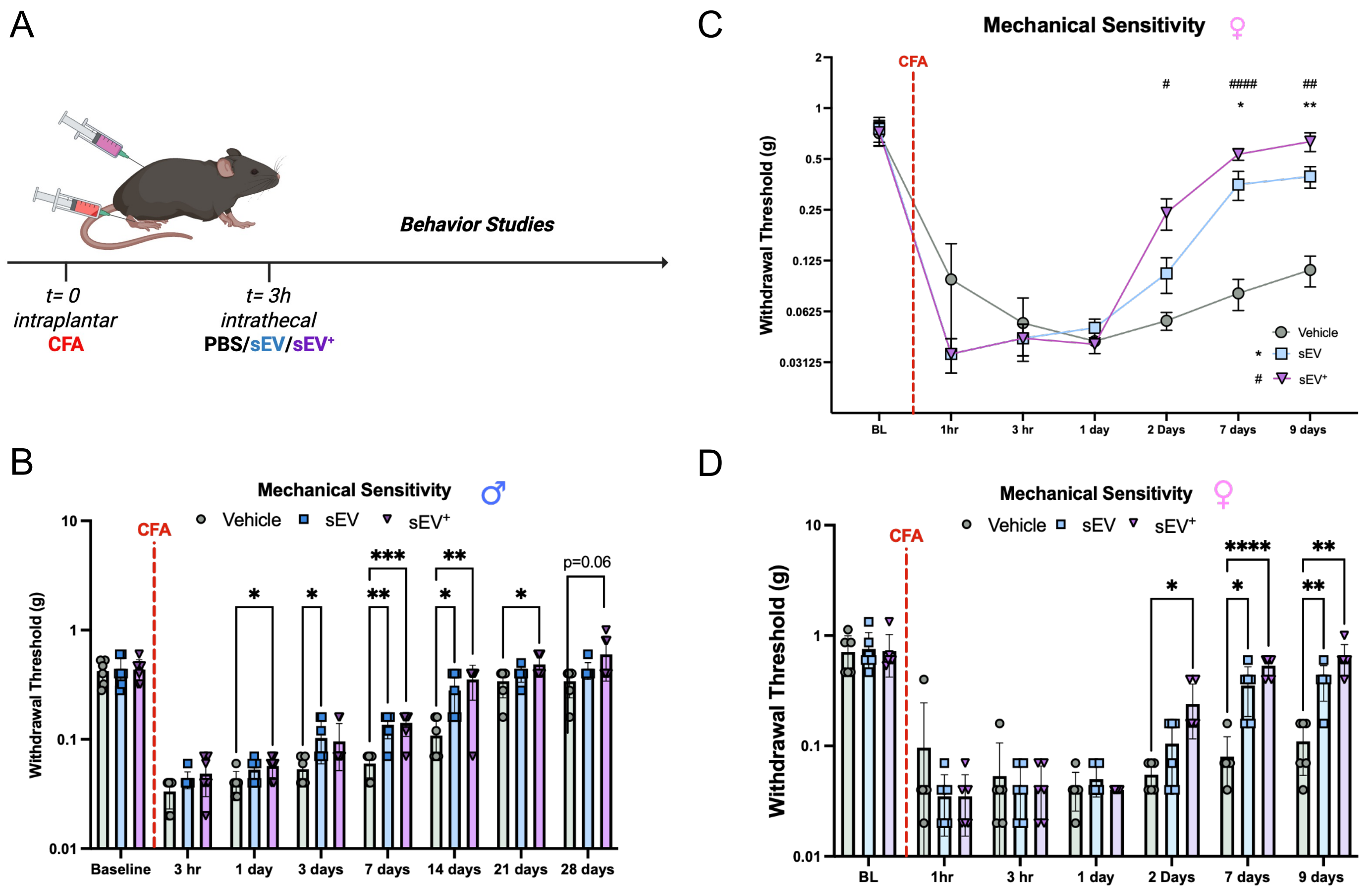

**Supplemental Figure 1. Intrathecal delivery of macrophage derived sEVs promotes early resolution of inflammatory pain and in male and female WT mice.** (**A**) Schematic showing timing of intraplantar CFA administration and intrathecal sEV administration (1 μg) as well as timepoints to assess mechanical sensitivity of the injected paw. (**B**) von Frey data from wild type male C57BL/6J mice in Figure 1E showing individual data points. Mean ± SEM (n=6-7), two-way repeated measures ANOVA with Dunnet's test for multiple comparisons, * $P<0.05$, ** $P<0.01$, *** $p<0.001$. sEV (derived from RAW 264.7 without LPS stimulation) shown in blue, and $sEV^+$ (derived from RAW 264.7 with LPS stimulation). Paw withdrawal threshold data from wild type female C57BL/6J mice shown as summary line graph (**C**) or as individual data points (**D**). Mean ± SEM (n=6), two-way repeated measures ANOVA with Dunnet's test for multiple comparisons. **C**) * PBS vs. sEV, # PBS vs. $sEV^+$. * $P<0.05$, ** $P<0.01$, **** $P<0.0001$. sEV shown in blue, and $sEV^+$ shown in purple.

### Supplemental Figure 2

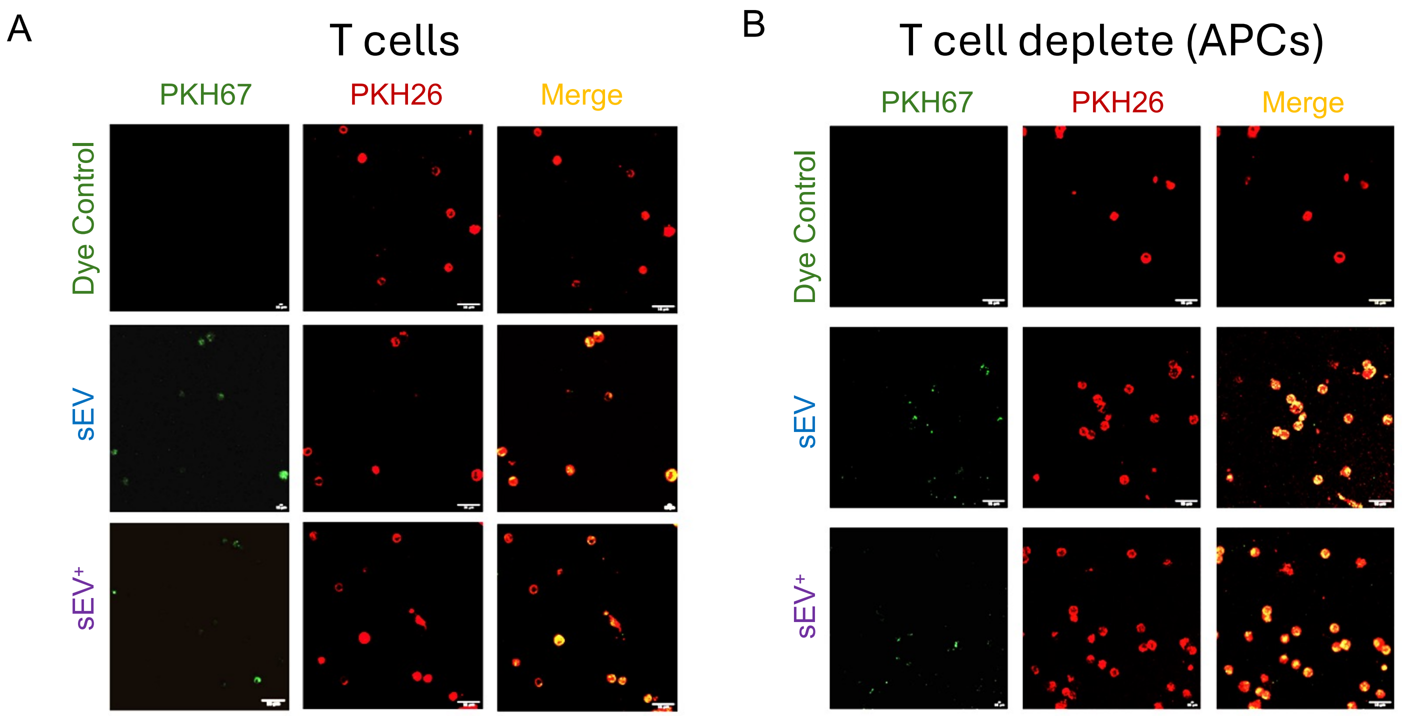

**Supplemental Figure 2. sEVs are taken up by both T cells and antigen presenting cells (APCs) *in* vitro.** Total T cells or T cell depleted (APCs) were labeled with lipophilic dye PKH26 and cultured with either PKH67 labeled sEVs (1 μg) or a dye control (remaining supernatant after washing during sEV labeling to account for residual dye and aggregates) for four hours at 37°C, fixed, and washed. Confocal images showing uptake of PKH67 labeled sEVs in both T cells (**A**) and APCs (**B**). Magnification 100x, scale 16 μm.

### Supplemental Figure 3

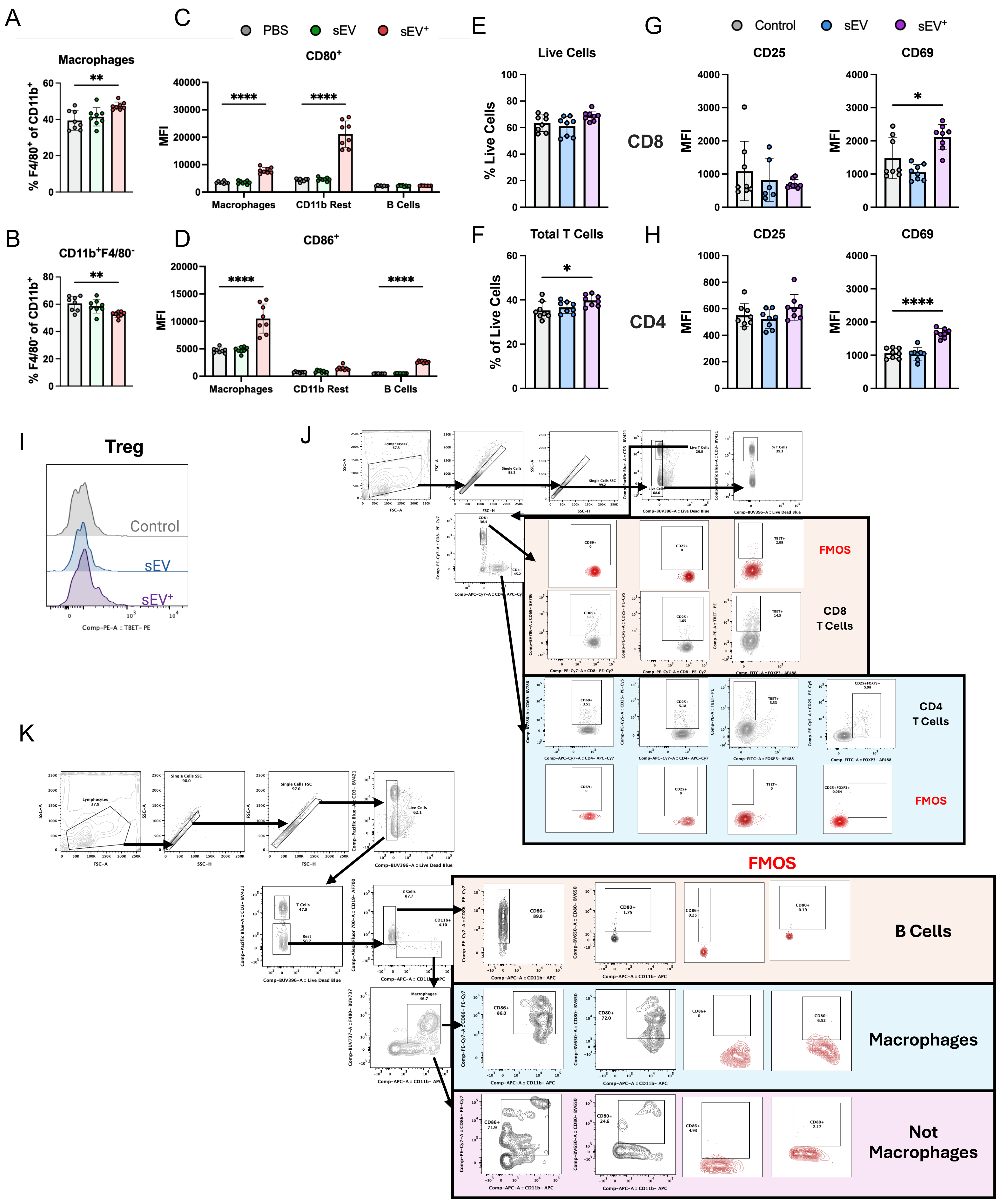

**Supplemental Figure 3. sEV$^+$ treatment alone increases APC and T cell activation in splenocyte cultures.** Indirect effects on APCs in splenocyte cultures (PBS- grey, sEV- green, sEV$^+$- red). (**A & B**) Percentages of macrophages (CD11b$^+$F4/80$^+$) and CD11b$^+$F4/80$^-$ of CD11b$^+$ cells. Mean ± SD (n=8), one-way ANOVA with Dunnet's test for multiple comparisons * $P$ <0.05, ** $P$ <0.01, **** $P$ <0.0001. (**C & D**) gMFI of positive populations in Figure 2C for CD80 and CD86 across APC populations. Two-way ANOVA with Dunnet's test for multiple comparisons * $P$ <0.05, *** $P$ <0.001, **** $P$ <0.0001. Indirect effects on T cells in splenocyte cultures 24 hours after treatment with 1 µg sEVs (PBS- grey, sEV- blue, sEV$^+$- purple). (**E & F**) Percentage of live cells and total T cells (CD3$^+$) in culture. (**G & H**) MFI of positive populations in Figure 2E & F of early (CD69) and late (CD25) activation markers. (**I**) Histogram of T-Bet expression in Treg from Figure 2G. Gating strategy for evaluation of T cells (**J**) and APC populations (**K**) in splenocyte cultures from Figure 2 and Supplemental Figure 3.

### Supplemental Figure 4

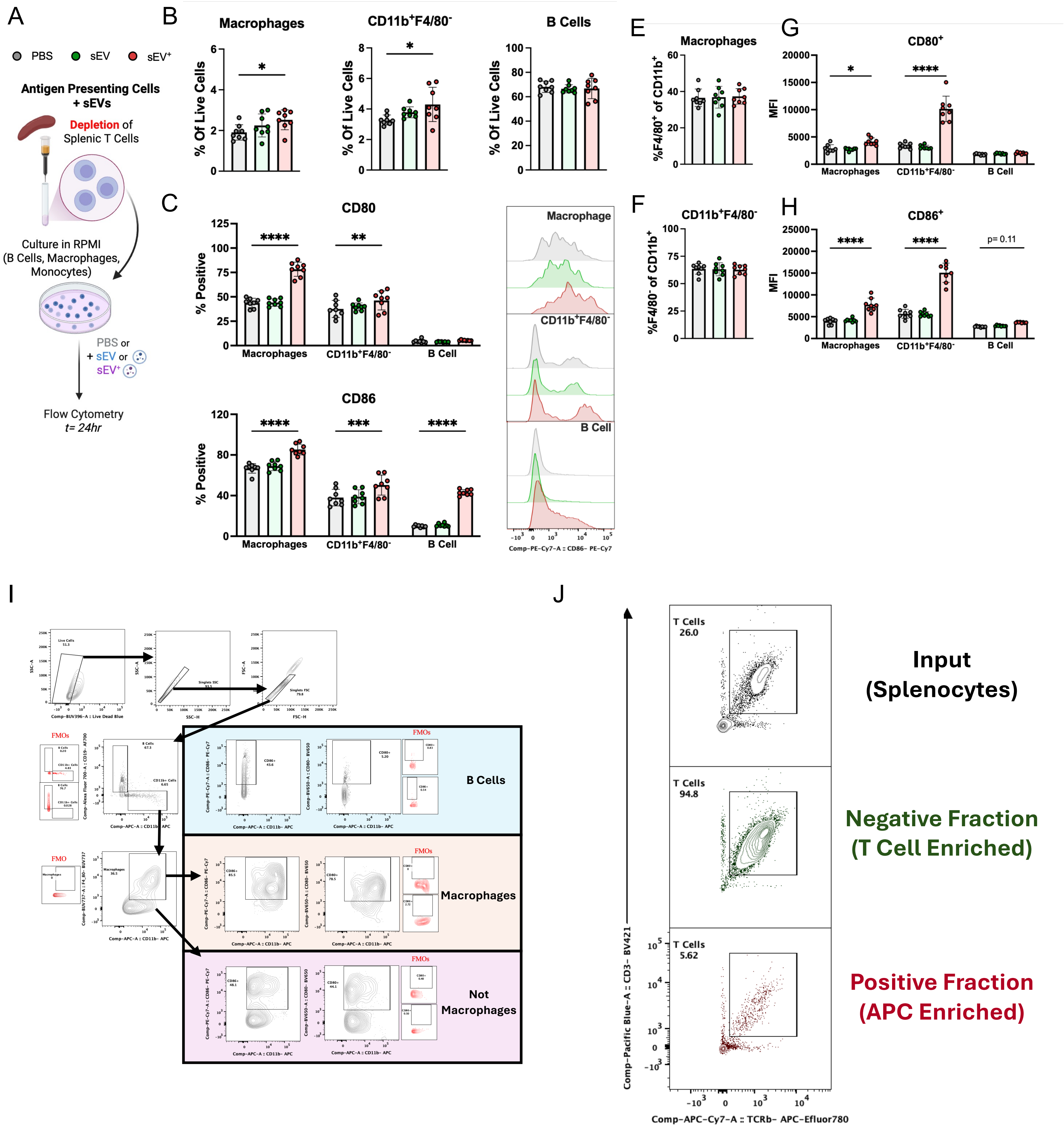

**Supplemental Figure 4. sEV$^+$ shows similar effects on APCs when treating T cell depleted cultures.** Direct effects on APCs in T cell depleted cultures 24 hours after treatment with 1 µg sEVs (PBS- grey, sEV- green, sEV$^+$- red). (**A**) Schematic for *in vitro* experiments using T cell depleted splenocytes cultured for 24 hours with or without sEVs (1 µg) to determine direct effects on APCS. (**B**) The effects of sEV treatment on APC percentages within the culture, including macrophages (CD11b$^+$F4/80$^+$), CD11b$^+$F4/80$^-$ (monocytes, dendritic cells, and neutrophils), and B cells (CD11b$^-$CD19$^+$). One-way ANOVA with Dunnet's test for multiple comparisons * p<0.05. (**C**) Expression of co-stimulation and activation markers CD80 (Top) and CD86 (Bottom) on APC populations in culture. Two-way ANOVA with Dunnet's test for multiple comparisons ** $P$<0.01, *** $P$<0.001, **** $P$<0.0001. (**D**) Concatenated histogram plots of APC populations from **C** showing increased CD86 expression on sEV$^+$ treated cells. (**E & F**) Percentages of macrophages (CD11b$^+$F4/80$^+$) and CD11b$^+$F4/80$^-$ of CD11b$^+$ cells. Mean ± SD *(n=8),* one-way ANOVA with Dunnet's test for multiple comparisons * $P$ <0.05, ** $P$ <0.01, **** $P$ <0.0001. (**G & H**) gMFI of positive populations in D for CD80 and CD86 across APC populations. Two-way ANOVA with Dunnet's test for multiple comparisons * $P$ <0.05, **** $P$ <0.0001. (**I**) Gating strategy for evaluation of APC populations in T cell deplete cultures. (**J**) Purity of MACS isolation of total T cell isolation showing input, negative fraction (total T cell enriched, CD3$^+$TCRβ$^+$) and positive fraction (APC Enriched).

### Supplemental Figure 5

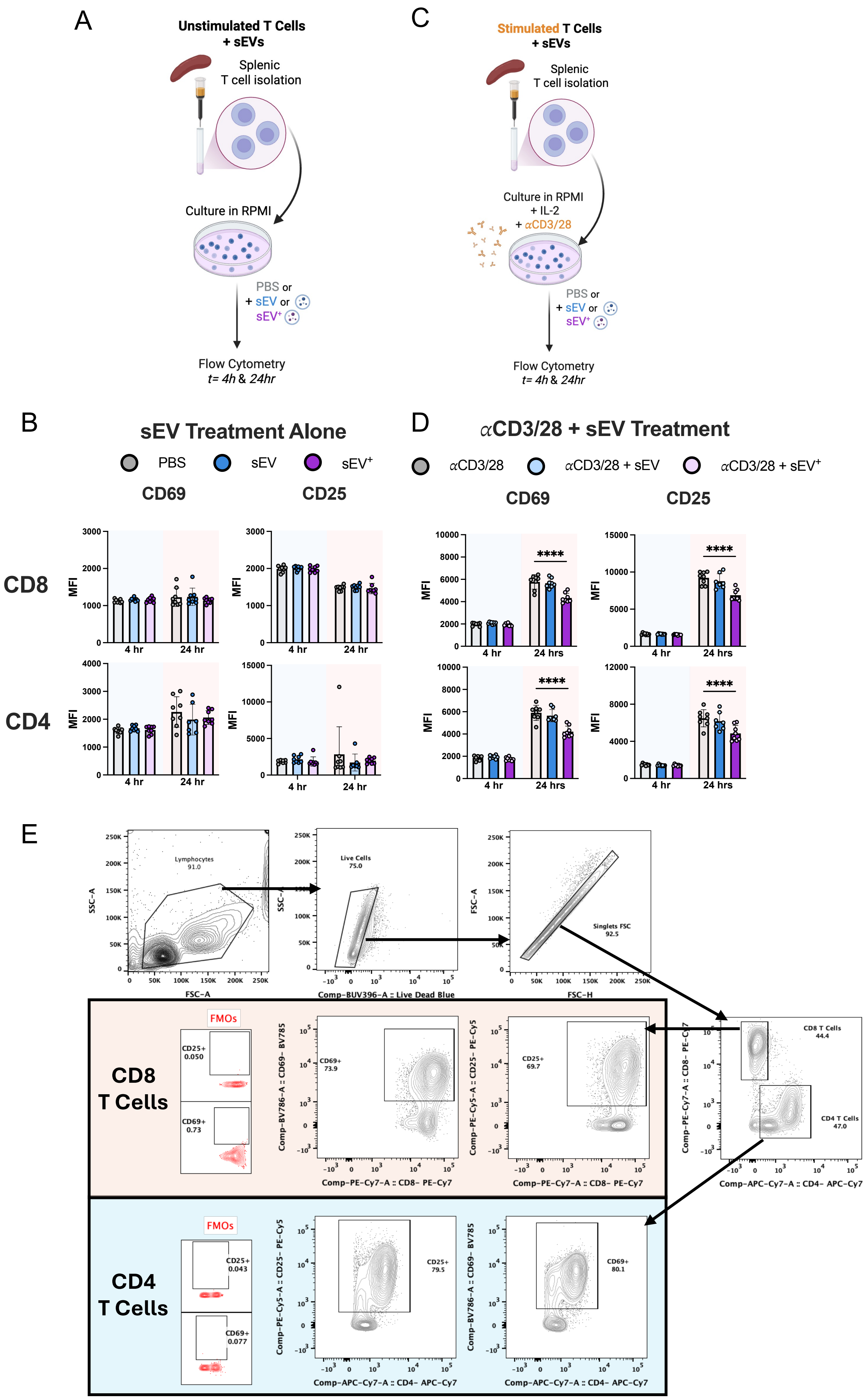

**Supplemental Figure 5. sEVs are capable of both directly activating and potentiating T cell activation.** (**A**) Schematic for *in vitro* experiments using splenic T cells isolated by negative magnetic selection and cultured for 4 or 24 hours with or without sEVs (1 μg) to determine direct effects. (**B**) The effects of sEV treatment alone on $CD8^+$ (Top) and CD4 (Bottom*)* expression *of* early (CD69) and late (CD25) activation markers as measured by gMFI of positive cell populations from Figure 3B. Mean ± SD (n=8), two-way ANOVA with Dunnet's test for multiple comparisons. (**C**) Schematic for *in vitro* experiments as in A but stimulated with $\alpha$-CD3/28 and IL-2 to examine if sEVs potentiate T cell activation. (**D**) The effects of sEV treatment of stimulated T cells on $CD8^+$ (Top) and $CD4^+$ (Bottom*)* expression *of* early (CD69) and late (CD25) activation markers as measured by gMFI of positive cell populations from Figure 3E. Mean ± SD (n=8), two-way ANOVA with Dunnet's test for multiple comparisons **** $P<0.0001$. (**E**) Gating strategy for experiments in Figure 3 and Supplemental Figure 5 with FMOS for terminal markers shown in red. sEV shown in blue, and $sEV^+$ shown in purple.

### Supplemental Figure 6

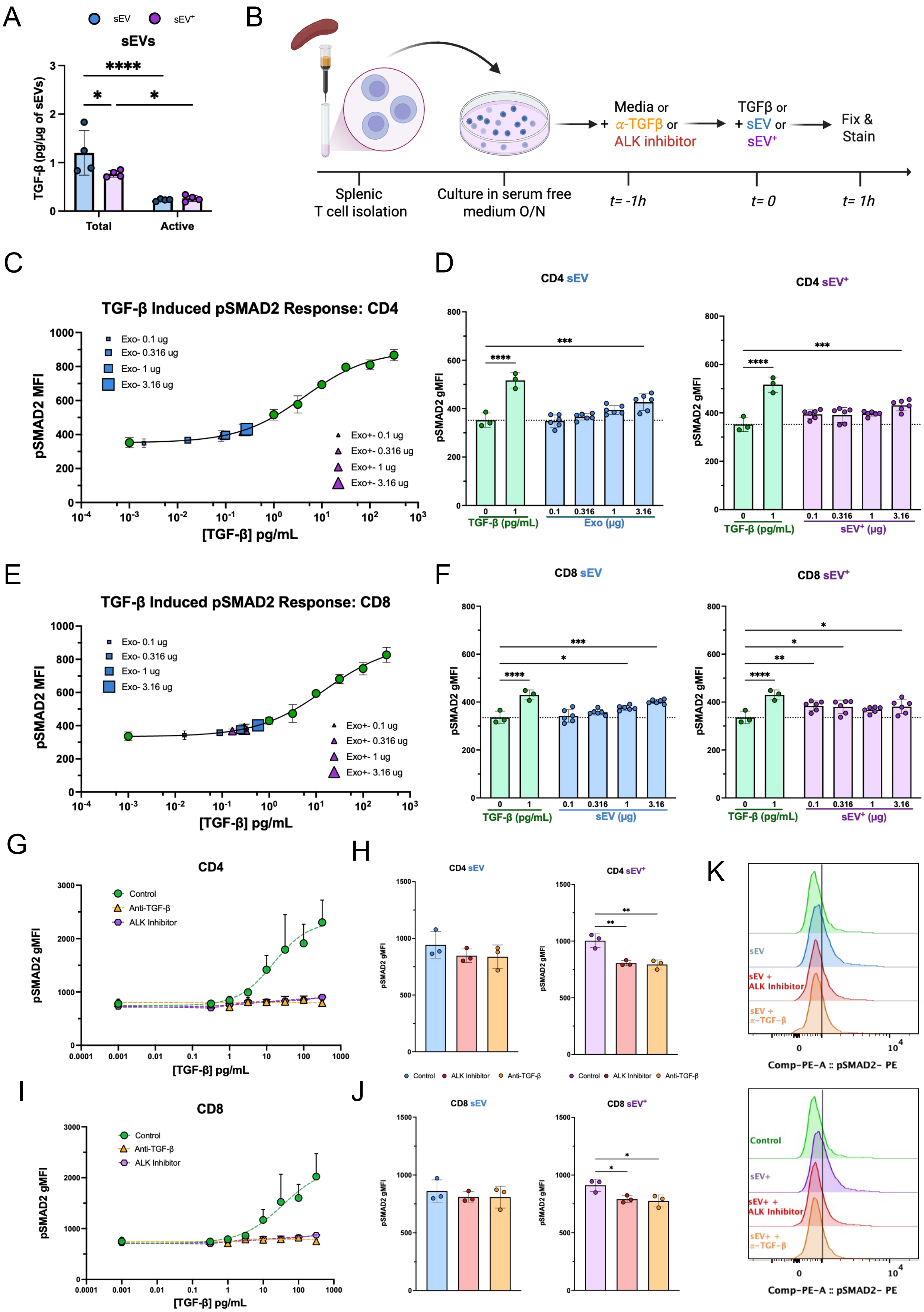

**Supplemental Figure 6. Macrophage derived sEVs contain TGF-β and induce pSMAD2 signaling.** (**A**) Amount of total and active TGF-β per µg of sEVs as measured by ELISA. Two-way ANOVA with Dunnet's test for multiple comparisons (n=4) * $P$ <0.05, **** $P$ <0.0001. (**B**) Schematic for experiments measuring pSMAD2 levels in T cells. Total T cells were isolated from splenocytes of naïve male mice via negative magnetic depletion and subsequently cultured in serum free media (as TGF-β exists at high concentrations in serum) for 16 hours over night. The subsequent day, in inhibitor experiments, cells were pre-treated with $\alpha$-TGF-β or ALK inhibitor for one-hour prior to the addition of sEVs or TGF-β as a positive control. At 1-hour post-treatment cells were immediately washed and methanol-fixed and stained for pSMAD2 signaling induced by TGF-β signaling in $CD4^+$ and $CD8^+$ cells by flow cytometry. (**C & E**) Representative dose response curves pSMAD2 signaling to TGF-β of in $CD4^+$ and $CD8^+$ T cells respectively, along with quantification of pSMAD2 MFI of sEV and $sEV^+$ treated cells (**D**) and (**F**) at different doses. 4PL-curve fits along with one-way ANOVA with Dunnet's test for multiple comparisons (n=3-6) * $P$ <0.05, ** $P$ <0.01, *** $P$ <0.001, **** $P$ <0.0001. Specificity of signal was tested by preincubation with either ALK inhibitor or $\alpha$-TGF-β prior to TGF-β treatment with respective curves in $CD4^+$ (**G**) and $CD8^+$ (**I**) T cells. (**H & J**) show specificity of sEV signal at a dose of 3.16 µg sEVs4PL-curve fits along with one-way ANOVA with Dunnet's test for multiple comparisons (n=3-6) * $P$ <0.05, ** $P$ <0.01, *** $P$ <0.001, **** $P$ <0.0001.

### Supplemental Figure 7

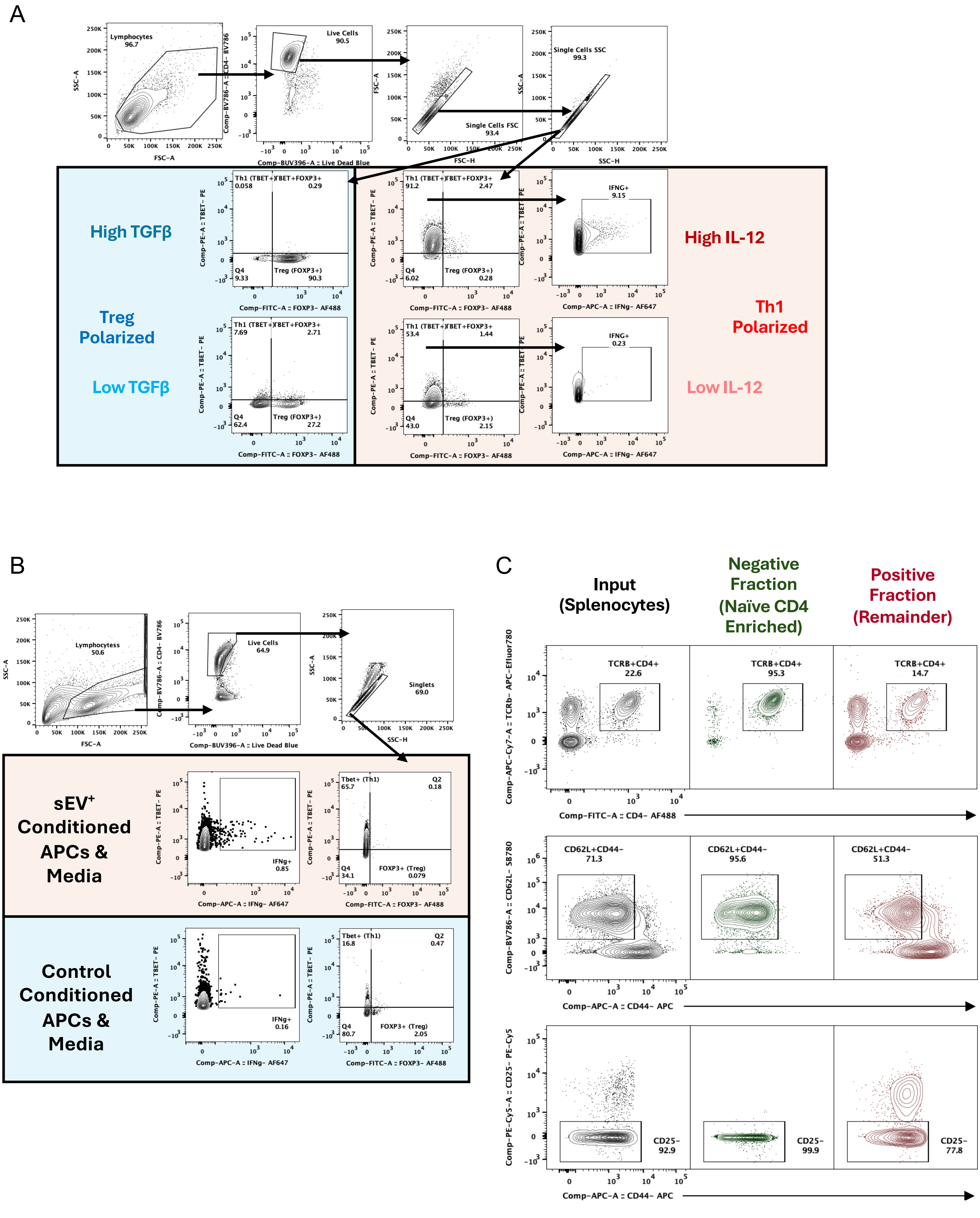

**Supplemental Figure 7. Gating strategies for T cell polarization experiments.** (**A**) Gating strategy and representative plots of $CD4^+$ naïve T cells treated with high and low concentrations of IL-12 and TGF-β from Figure 4A-E. (**B**) Gating strategy and representative plots of $CD4^+$ naïve T cells treated conditioned APCs and media Figure 4F-K. (**C**) Purity of MACS isolation of CD4 Naïve T cells showing input, negative fraction (Naïve CD4 enriched, $CD4^+TCR\beta^+CD62L^+CD44^-CD25^-$), and positive fraction (remainder).

### Supplemental Figure 8

A

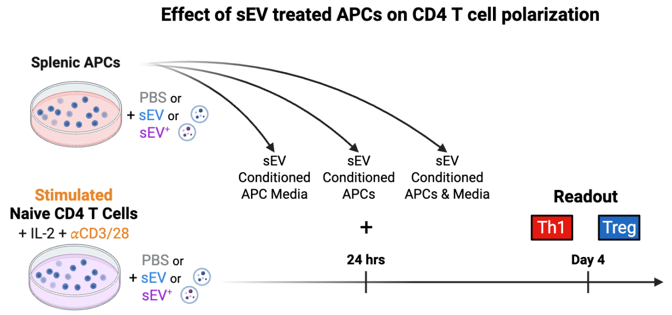


B

**Treg**

| Control | sEV T Cells + APC Media | sEV T Cells + APCs | sEV T Cells + APCs & Media |
|---|---|---|---|
| sEV T Cells | ns | **** | **** |
| sEV T Cells + APC Media | | **** | **** |
| sEV T Cells + APCs | | | ns |

| sEV | sEV T Cells + APC Media | sEV T Cells + APCs | sEV T Cells + APCs & Media |
|---|---|---|---|
| sEV T Cells | ns | *** | *** |
| sEV T Cells + APC Media | | ** | ns |
| sEV T Cells + APCs | | | ns |

| sEV+ | sEV T Cells + APC Media | sEV T Cells + APCs | sEV T Cells + APCs & Media |
|---|---|---|---|
| sEV T Cells | **** | **** | **** |
| sEV T Cells + APC Media | | ns | ns |
| sEV T Cells + APCs | | | ns |

**Th1**

| Control | sEV T Cells + APC Media | sEV T Cells + APCs | sEV T Cells + APCs & Media |
|---|---|---|---|
| sEV T Cells | ns | **** | **** |
| sEV T Cells + APC Media | | **** | **** |
| sEV T Cells + APCs | | | * |

| sEV | sEV T Cells + APC Media | sEV T Cells + APCs | sEV T Cells + APCs & Media |
|---|---|---|---|
| sEV T Cells | ns | *** | ns |
| sEV T Cells + APC Media | | *** | ns |
| sEV T Cells + APCs | | | *** |

| sEV+ | sEV T Cells + APC Media | sEV T Cells + APCs | sEV T Cells + APCs & Media |
|---|---|---|---|
| sEV T Cells | ** | **** | **** |
| sEV T Cells + APC Media | | **** | **** |
| sEV T Cells + APCs | | | ns |

**Th1- IFNγ+**

| Control | sEV T Cells + APC Media | sEV T Cells + APCs | sEV T Cells + APCs & Media |
|---|---|---|---|
| sEV T Cells | ns | ns | ns |
| sEV T Cells + APC Media | | ns | ns |
| sEV T Cells + APCs | | | ns |

| sEV | sEV T Cells + APC Media | sEV T Cells + APCs | sEV T Cells + APCs & Media |
|---|---|---|---|
| sEV T Cells | ns | ns | * |
| sEV T Cells + APC Media | | ns | * |
| sEV T Cells + APCs | | | ns |

| sEV+ | sEV T Cells + APC Media | sEV T Cells + APCs | sEV T Cells + APCs & Media |
|---|---|---|---|
| sEV T Cells | ns | **** | **** |
| sEV T Cells + APC Media | | **** | **** |
| sEV T Cells + APCs | | | ns |

**Supplemental Figure 8. sEV$^+$ suppresses Treg and promotes Th1 polarization through both direct and indirect effects.** (**A**) Schematic for *in vitro* polarization experiments in which naïve CD4$^+$ T cells isolated by negative magnetic selection from lymph nodes and spleens of naïve mice were stimulated with $\alpha$-CD3/28 and IL-2, with or without sEVs for 24 hours. Additionally, APCs (T cell deplete) were also treated with or without sEVs for 24 hours. After 24 hours APC cultures were spun down and separated into conditioned media and APCs. Conditioned media, APCs, or both were supplemented to T cell cultures at 24 hours in a matched fashion for APC/T cell cultures (PBS/PBS, sEV/sEV, sEV$^+$/sEV$^+$). Cells were removed from stimulation on day 2 and supplemented with IL-2 each day until day 4 when they were restimulated ($\alpha$-CD3/28) and analyzed. (**B**) Comparisons within treatment condition and between groups from **Figure 4G-I**. Mean ± SD (n=8), two-way ANOVA with Dunnet's test for multiple comparisons * $P$ <0.05, ** $P$ <0.01, *** $P$ <0.001, **** $P$ <0.0001.

### Supplemental Figure 9

A

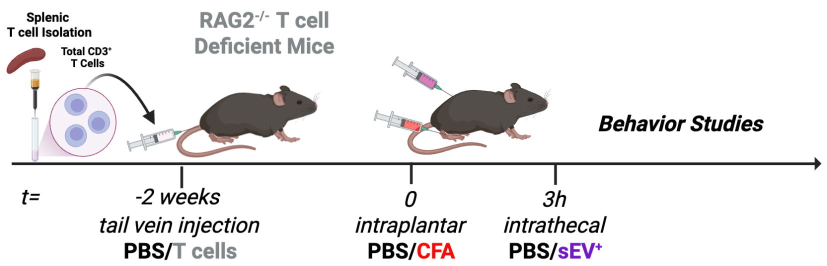


B

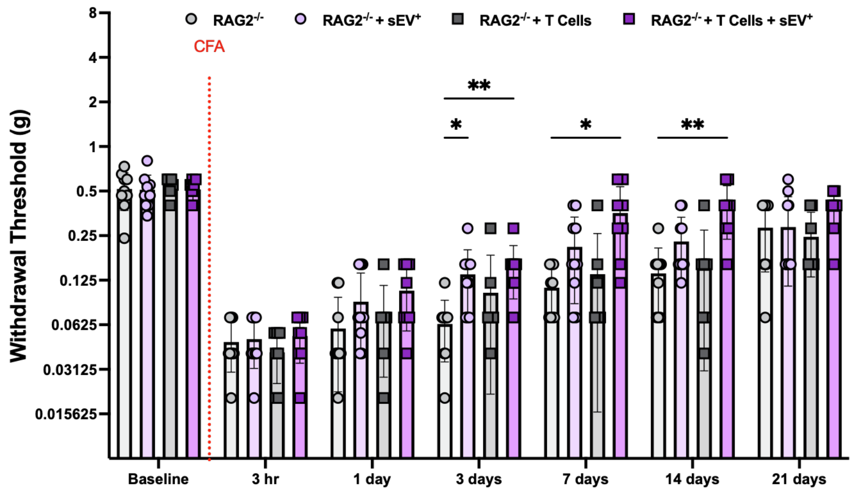

**Supplemental Figure 9. Late-phase inflammatory pain resolution mediated by intrathecal delivery of sEV$^+$ requires T cells.** (**A**) Schematic showing design for adoptive transfer of total CD3$^+$ T cells isolated from WT C57BL/6J mice by negative magnetic selection of splenocytes and injected into the tail vein of T cell deficient *Rag2*$^{-/-}$ mice. Two weeks after adoptive transfer, mice were subjected to intraplantar CFA and intrathecal sEV administration (1 μg) and assessed for mechanical sensitivity of the injected paw via von Frey filaments for up to 3 weeks. (**B**) von Frey data evaluating mechanical sensitivity of *Rag2*$^{-/-}$ mice with or without T cell transfer and treatment with sEV$^+$ from Figure 6D showing individual data points. Mean ± SEM (n=7-10), two-way repeated measures ANOVA with Dunnet's test for multiple comparisons * $P$ <0.05, ** $P$ <0.01.

### Supplemental Figure 10

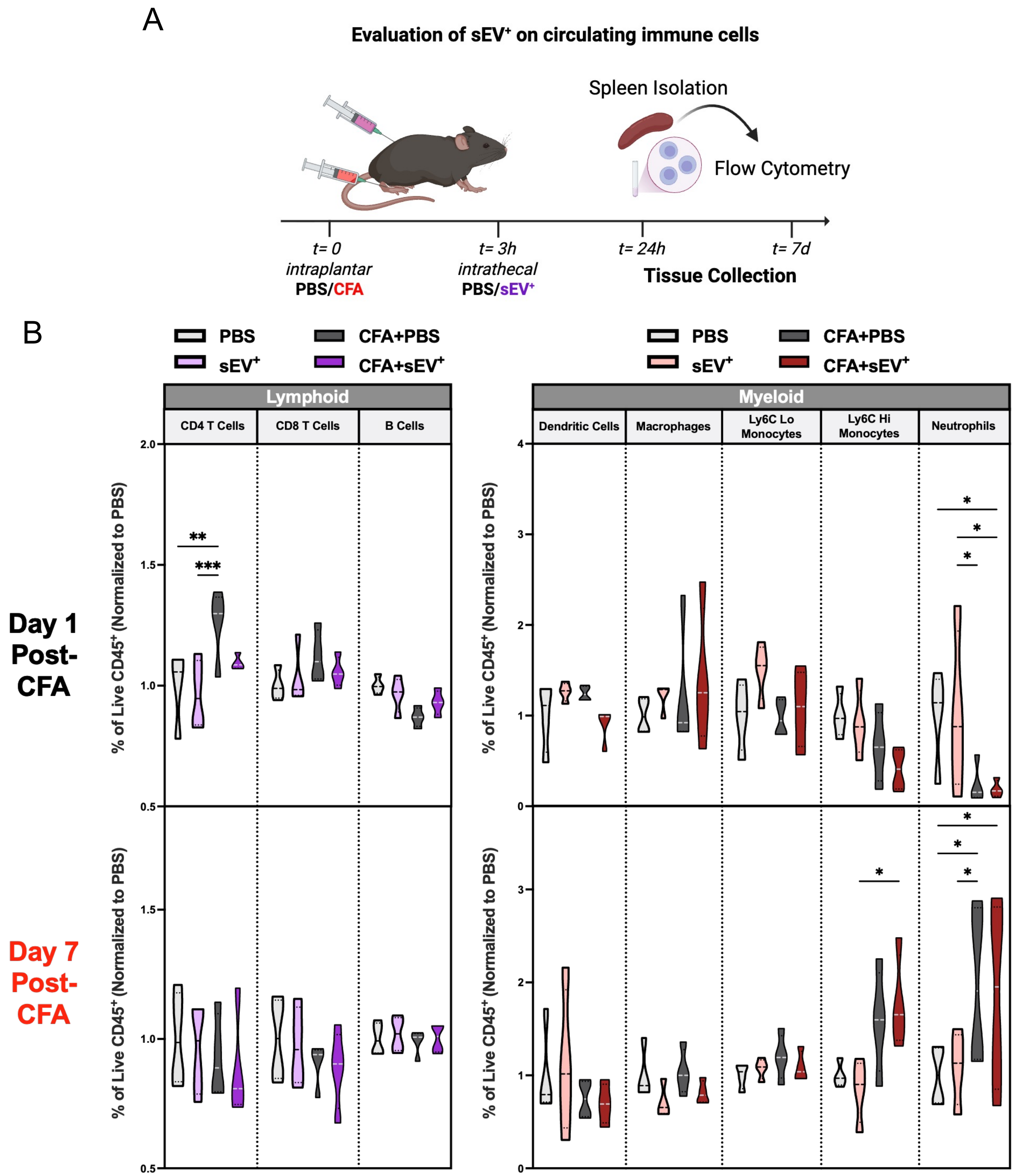

Supplemental Figure 10 cont.

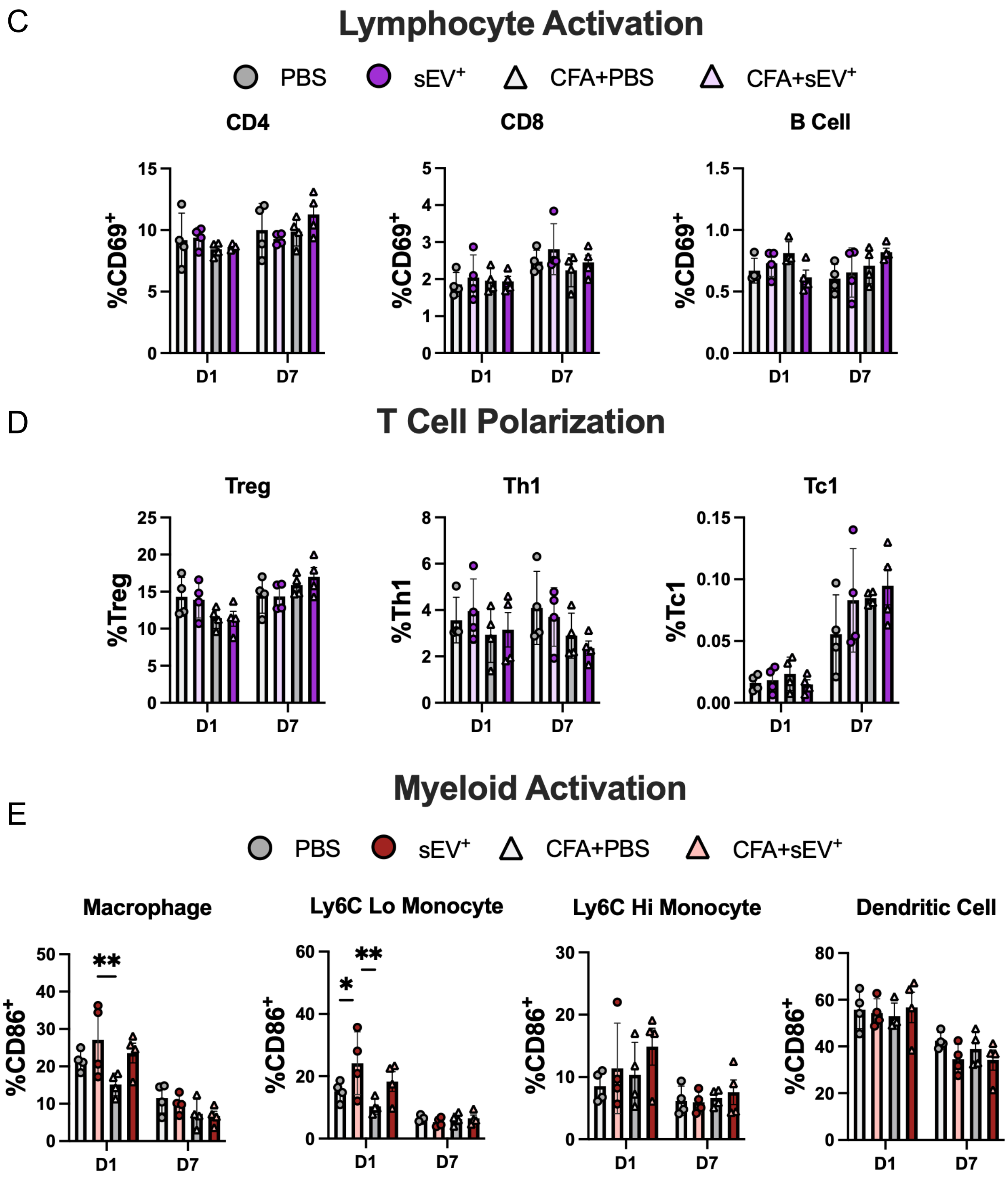

Supplemental Figure 10 cont.

F

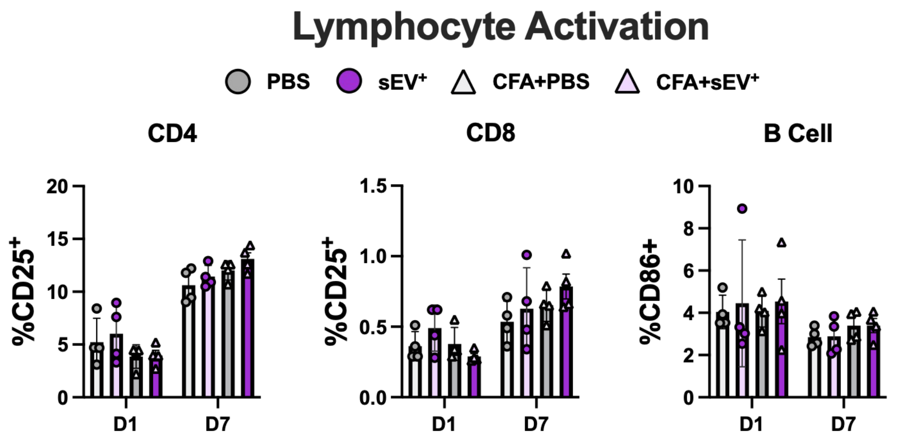


G

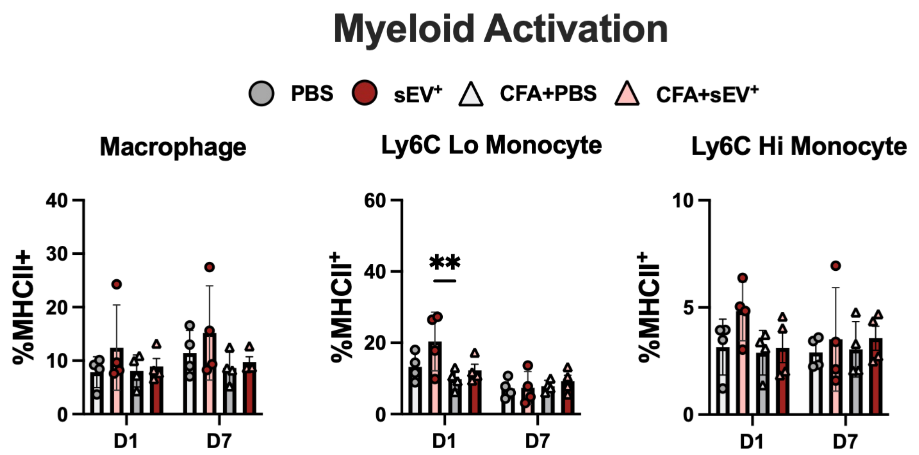


H

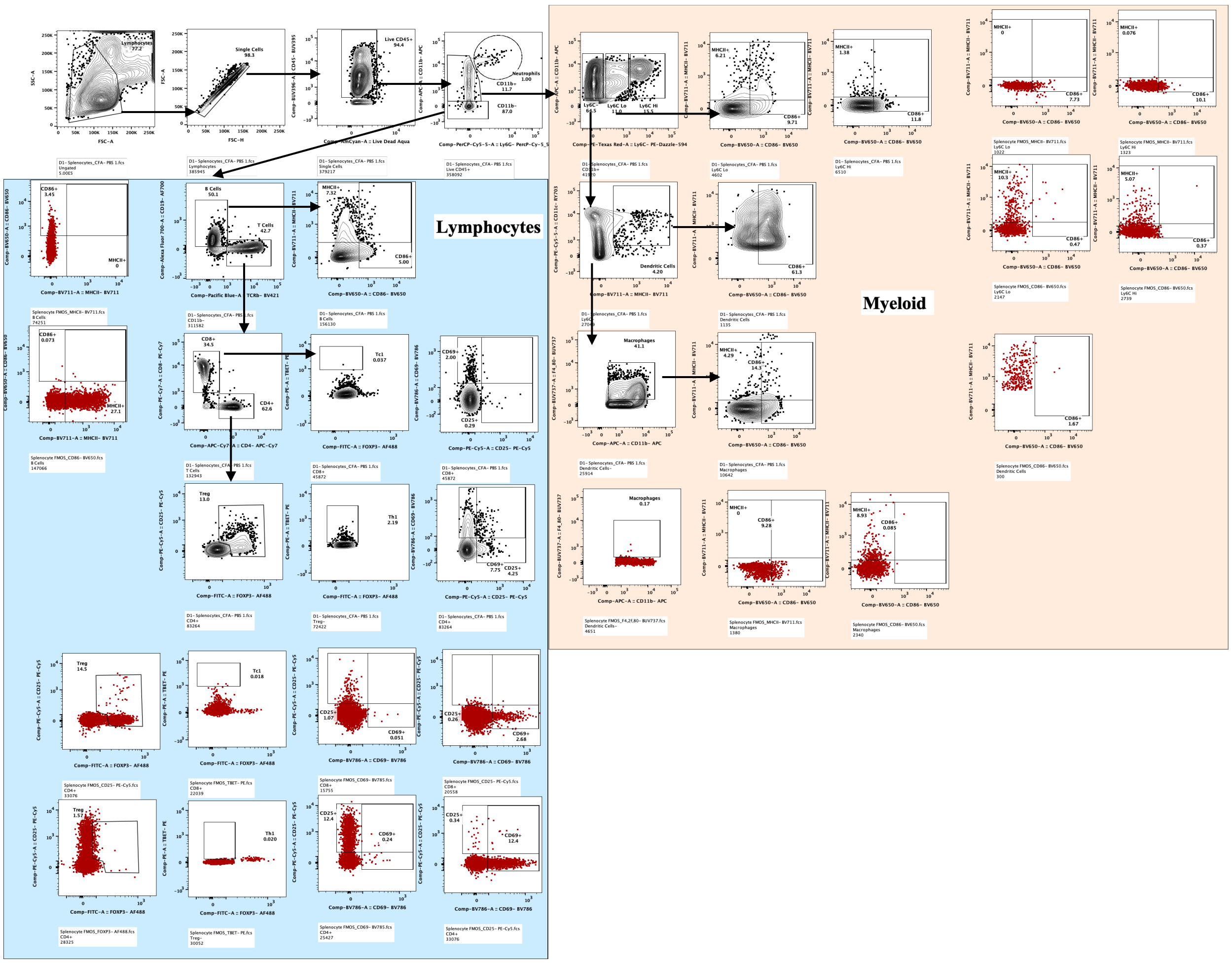

**Supplemental Figure 10. Intrathecal delivery of $sEV^+$ has minimal effects on splenic immune populations.** (**A**) Schematic showing design to evaluate splenic populations at 1 and 7 days after CFA/$sEV^+$ administration via flow cytometry. (**B**) Truncated violin plots showing the percentage of lymphoid (left) (PBS- light grey, $sEV^+$- light purple, CFA+PBS- dark grey, CFA+$sEV^+$- dark purple) and myeloid (right) (PBS- light grey, $sEV^+$- light red, CFA+PBS- dark grey, CFA+$sEV^+$- dark red) cell populations in the spleen at day 1 (top) and day 7 (bottom) post CFA/$sEV^+$ administration. Cell percentages were normalized by cell type to PBS control. Lymphoid and myeloid were separated due to differences in variance as lymphoid represent >90% of cells in the spleen while myeloid is <10% (Supplemental Figure 11B); (n=4) two-way ANOVA with Dunnet's test for multiple comparisons. * $P$<0.05, ** $P$<0.01, *** $P$<0.001. (**C**) Lymphocyte activation as measured by expression of early activation marker CD69 across lymphocyte populations. (**D**) T cell polarization as measured by percentage of Treg ($CD4^+FOXP3^+$), Th1 ($CD4^+$T-$Bet^+$), and Tc1 ($CD8^+$T-$Bet^+$). (**E**) Myeloid activation as measured by expression of co-stimulatory signal CD86 across macrophages ($CD11b^+Ly6G^-Ly6C^-$ $F4/80^+$), Ly6C Lo and Hi monocytes ($CD11b^+Ly6G^-Ly6C^{Lo/Hi}$), and dendritic cells ($CD11b^+Ly6G^-Ly6C^-CD11c^+MHCII^+$). Alternate activation markers for splenic lymphocyte (PBS- light grey, $sEV^+$- light purple, CFA+PBS- dark grey, CFA+$sEV^+$- dark purple) and myeloid (PBS- light grey, $sEV^+$- light red, CFA+PBS- dark grey, CFA+$sEV^+$- dark red) from A-E. (**F**) Lymphocyte activation as measured by expression of late activation marker CD25 (T cell) and alternate co-stimulatory molecule CD86 (B cell) across lymphocyte populations. (**G**) Myeloid activation as measured by expression of co-stimulatory signal CD86 across macrophages ($CD11b^+Ly6G^-Ly6C^-F4/80^+$) and Ly6C Lo and Hi monocytes ($CD11b^+Ly6G^-$ $Ly6C^{Lo/Hi}$). (**H**) Gating strategy for experiments from Supplemental Figure 10 & Figure 8 showing representative populations from the spleen with representative FMOs in red for terminal marker expression.

### Supplemental Figure 11

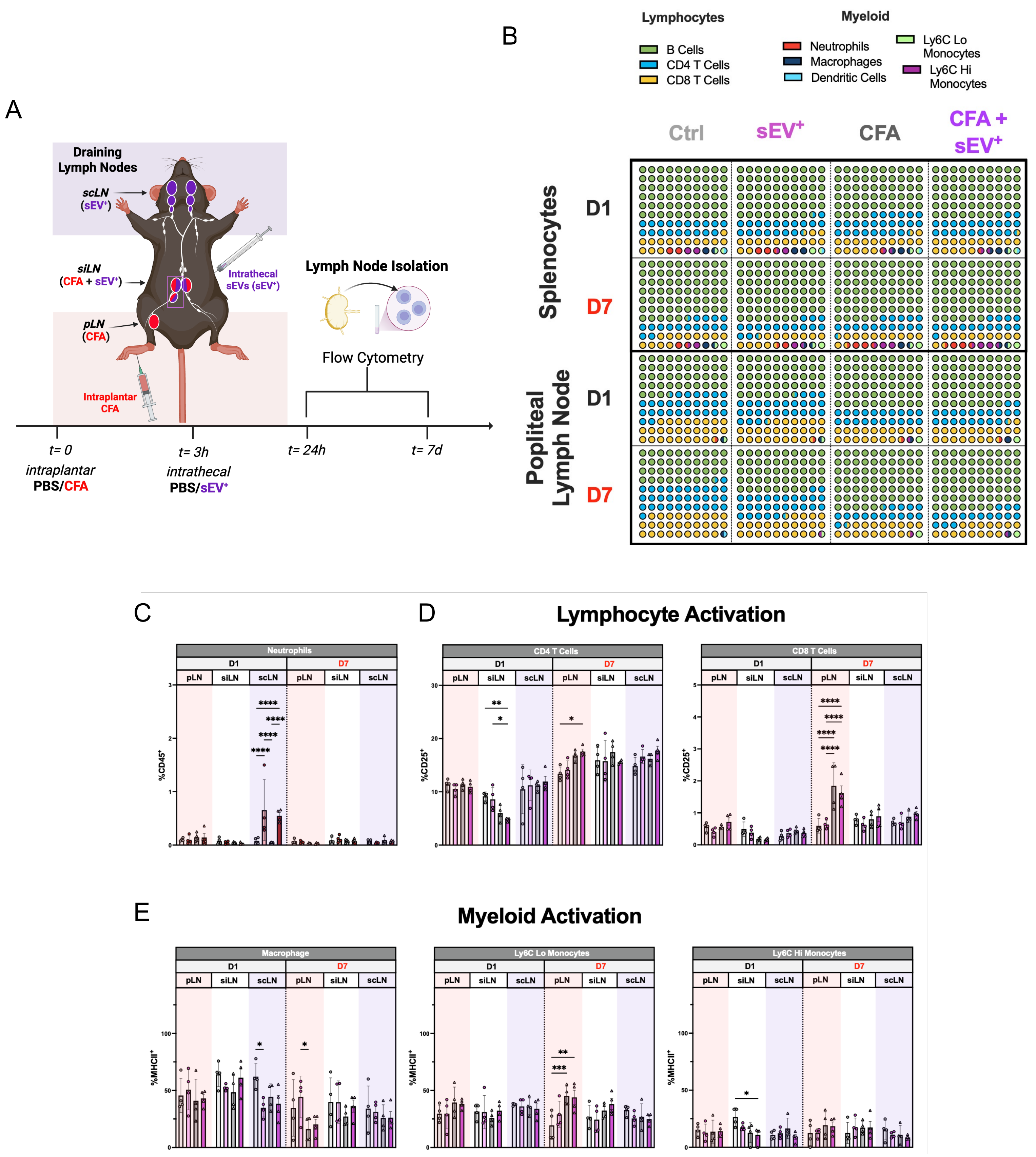

**Supplemental Figure 11. Intrathecal delivery of $sEV^+$ strongly activates immune populations in the cervical draining lymph nodes.** (**A**) Schematic showing design to evaluate differences in immune cell activation of immune cell populations in draining lymph nodes. Draining lymph nodes (LN) were assessed for intraplantar injection (popliteal (p)LN, sacral/internal iliac (si)LN) and intrathecal injection (superior cervical (sc)LN, and siLN) where pLN is unique to CFA injection (light red background), scLN is unique to $sEV^+$ injection (light purple background) and the siLN represents a potential intersecting draining site (overlap or white background in figures). Immune cell populations from draining lymph nodes were assessed at 1 and 7 days after CFA/$sEV^+$ injection to assess lymphoid (PBS- light grey, $sEV^+$- light purple, CFA+PBS- dark grey, CFA+$sEV^+$- dark purple) and myeloid (PBS- light grey, $sEV^+$- light red, CFA+PBS- dark grey, CFA+$sEV^+$- dark red) populations via flow cytometry. (**B**) Parts of whole graphs showing the distribution of different cell populations across spleen and a representative lymph node (popliteal), where each circle represents 1%. (**C**) Neutrophils measured in lymph nodes at day 1 (Left) and day 7 (Right), which were analyzed separately due to the low % typically seen in LNs (<0.1%). (**D**) Alternate lymphocyte activation marker CD25 across T cell populations from Figure 7E. **(E**) Alternate myeloid activation and antigen presentation marker MHCII across macrophages ($CD11b^+Ly6G^-Ly6C^-F4/80^+$), and Ly6C Lo and Hi monocytes ($CD11b^+Ly6G^-Ly6C^{Lo/Hi}$) from Figure 7F. (n=4) two-way ANOVA with Dunnet's test for multiple comparisons * $P$ <0.05, ** $P$ <0.01, *** $P$ <0.001, **** $P$ <0.0001.
